# Waterlogging stress induces antioxidant defense responses and aerenchyma formation and alters metabolisms of banana plants

**DOI:** 10.1101/2022.06.21.496926

**Authors:** Ee Yang Teoh, Chee How Teo, Nadiya Akmal Baharum, Teen-Lee Pua, Boon Chin Tan

**Affiliations:** Centre for Research in Biotechnology for Agriculture (CEBAR), Universiti Malaya, 50603 Kuala Lumpur, Malaysia; Department of Cell and Molecular Biology, Faculty of Biotechnology and Biomolecular Sciences, Universiti Putra Malaysia, 43400 UPM Serdang, Selangor Darul Ehsan, Malaysia; Topplant Laboratories Sdn. Bhd., Jalan Ulu Beranang, 71750, Lenggeng, Negeri Sembilan, Malaysia

**Keywords:** Abiotic stress, Banana, Crop improvement, Waterlogging, Transcriptomics

## Abstract

Flooding caused or exacerbated by climate change has threatened plant growth and food production worldwide. The lack of knowledge on how crops respond and adapt to flooding stress imposes a major barrier to enhancing their productivity. Hence, understanding the flooding-responsive mechanisms of crops is indispensable for developing new flooding-tolerant varieties. Here, we examined the banana *(Musa acuminata* cv. Berangan) responses to soil waterlogging for 1, 3, 5, 7, 14, and 24 days. After waterlogging stress, banana root samples were analyzed for their molecular and biochemical changes. We found that waterlogging treatment induced the formation of adventitious roots and aerenchyma with conspicuous gas spaces. In addition, the antioxidant activities, hydrogen peroxide and malondialdehyde contents of the waterlogged bananas increased in response to waterlogging stress. To assess the initial response of bananas toward waterlogging stress, we analyzed the transcriptome changes of banana roots. A total of 3,508 unigenes were differentially expressed under 1-day waterlogging conditions. These unigenes comprise abiotic stress-related transcription factors, such as ethylene response factors, basic helix-loop-helix, myeloblastosis, plant signal transduction, and carbohydrate metabolisms. The findings of the study provide insight into the complex molecular events of bananas in response to waterlogging stress, which could later help develop waterlogging resilient crops for the future climate.

## 1. Introduction

Flooding is one of the most significant threats to food production and economic growth worldwide. It affects 17 million km^2^ of land surface annually [1], causing severe damage to agricultural crop production. The frequency of flooding events is expected to increase in the near future, especially in Southern Asia countries [2], due to increasingly erratic rainfall patterns exacerbated by climate change [3]. For example, in Malaysia, the flooding event in December 2006 amounted to USD 18.9 million of agriculture loss and damage, affecting the arable lands and farmers [4]. Similarly, the flooding event in the Mississippi River and Midwest in 2019 resulted in a cumulative loss of USD 6.9 billion, mainly affecting agriculture [5]. These occurrences encourage scientific initiatives to develop flood-tolerant varieties to mitigate agricultural losses and improve global crop production.

Waterlogging is part of the flooding stress, where the soil is oversaturated with water, and only plant roots are surrounded by water. Oxygen partial pressure and gas diffusion in waterlogged soil gradually decline, disrupting plant root function and normal cellular processes and metabolisms [6]. Consequently, the nutrient uptake by plants dramatically decreases, leading to plant starvation [7]. Several studies have shown that waterlogging is a significant yield-limiting factor for crop yield and growth [8,9]. For instance, waterlogging reduced the yield of corn by about 4.7% for each day of waterlogging for up to seven days compared to well-watered corn [10]. Therefore, it is crucial to understand the plant molecular mechanisms in response to flooding stress to mitigate its effects on plants.

Plants have developed multiple and interrelated signaling pathways to modulate stress-responsive genes, leading to morphological, physiological, and metabolic changes [11]. At the morphological level, plant growth, biomass, and yield decreased under waterlogged conditions [12]. In addition, root architecture for the waterlogged-tolerant plant species is usually altered under waterlogging conditions. For instance, adventitious roots and aerenchyma were developed in maize in response to waterlogging [13]. The formation of adventitious roots and aerenchyma could help mitigate oxidative stress and promote oxygen movement to roots under hypoxic stress [14]. Besides changing the root system, plants reduce their photosynthetic capacity and increase antioxidant enzyme activities when exposed to stress [15]. At the physiological and biochemical levels, waterlogging stress induces stomatal closure, osmolyte accumulation, and reactive oxygen species (ROS) scavenging mechanism [16]. In the presence of ethylene, waterlogging stress enhances respiratory burst oxidase homolog (RBOH) expression, leading to the increase of ROS and hydrogen peroxide (H2O2) levels [17]. At the molecular level, genes and proteins involved in hormonal signal pathways, ROS production, and carbohydrate and energy metabolisms have been reported to play significant roles in waterlogging stress responses through the N-end rule pathway [18]. These findings show that plants adopt different strategies to defend themselves from waterlogging stress.

Recently, much progress has been made to decipher the plant responsive mechanisms against flooding stress [19,20]. However, although the previous findings contribute to our understanding of the adverse effects of flooding stress on plants, how crops respond and adapt to flooding stress remains largely unknown, imposing a major barrier to enhancing their productivity. Hence, understanding the flooding-responsive mechanisms of crops is indispensable for developing new flooding-tolerance varieties.

Banana is a commercially important cash crop as its nutritional value is higher than other tropical fruits [21]. It ranked among the top ten crops in terms of yield and calories produced, with about 119 million tons produced globally in 2020 [22]. However, global extreme precipitation events are a major constraint for banana yield and negatively affect its productivity [23]. Furthermore, the predicted rise in the earth’s global temperature is likely to aggravate flooding effects, fostering a global decline in banana yield. Therefore, improving banana production and the quality of this economically valuable fruit is vital.

In this study, we examined the growth and global gene expression of banana plants *(M. acuminata* cv. Berangan) in response to waterlogging stress to better understand the mechanism involved in the waterlogging tolerance. We determined the morphological and biochemical changes of banana plants at different time points of waterlogging treatment. In addition, several genes involved in waterlogging signaling and regulation in the early stage of waterlogging were also identified using transcriptomics and bioinformatics approaches.

## 2. Results

### 2.1. Waterlogging Stress Influences Banana Growth

The growth of well-watered and waterlogged banana plants was measured at 1, 3, 5, 7, 14, and 24 days (Figure 1). Waterlogging stress impacted the banana root length, producing the highest root length of 272.4 cm after 24 days of waterlogging (Figure 1A). The root-to-shoot (R/S) dry weight (DW) for well-watered samples increased from 0.18 g g^-1^ on day 1 to 0.26 g g^-1^ on day 24 (Figure 1B). In contrast, a slight increase in the R/S DW ratio for waterlogged samples was observed after 7 days, from 0.15 g g^-1^ to 0.26 g g^-1^ on day 24 (Figure 1B). The waterlogged plants generally had a larger leaf area than well-watered plants. The leaf area differences were highest on day 3 for well-watered plants (237.52 cm^2^) and waterlogged bananas (459.3 cm^2^) (Figure 1C). However, the differences between well-watered and waterlogged samples were not significant. The leaves of the waterlogged banana showed yellowing after 24 days of waterlogging (Supplementary Figure 1). The relative water content (RWC) for both treatments was similar throughout the experiments, ranging between 82 % and 95 % (Figure 1D).

**Figure 1.**
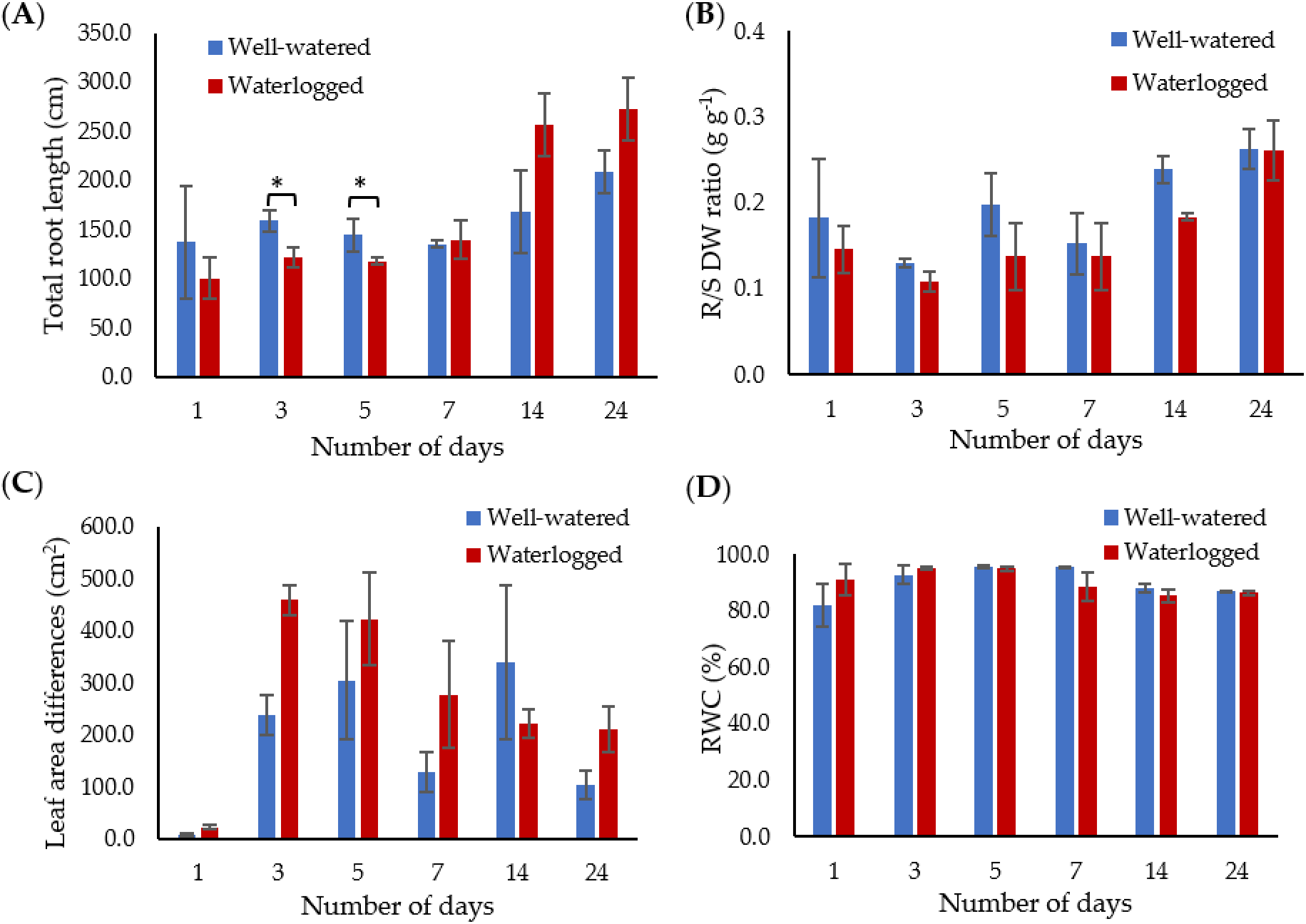
The growth of well-watered and waterlogged banana plants was measured in terms of (**A**) total root length, (**B**) root-to-shoot (R/S) dry weight (DW) ratio, and (**C**) leaf area differences. (**D**) The effect of waterlogging on relative water content (RWC) was assessed between well-watered and waterlogged leaf samples. Data are presented as means ± standard error from three independent biological replicates. Asterisk (*) indicates that the treatment was significantly different *(p* < 0.05).

### 2.2. Waterlogging Induces Adventitious Roots and Aerenchyma Formation

The waterlogged banana plants started to produce adventitious roots after 3 days of waterlogging, with the highest number of adventitious roots (10 roots) recorded after 24 days of waterlogging (Figure 2A). In contrast, the well-watered plants did not grow adventitious roots throughout the experiment (Figure 2A).

**Figure 2.**
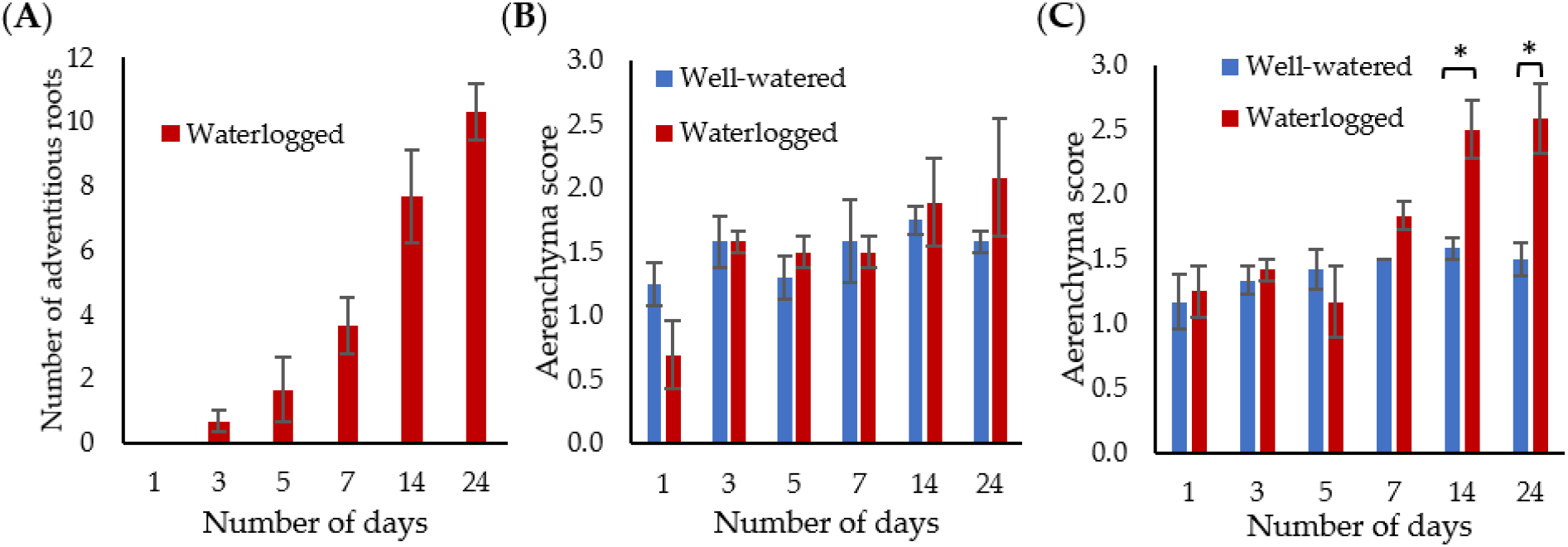
Adventitious root and aerenchyma formation. (**A**) The number of adventitious roots in well-watered and waterlogged banana plants. The mean aerenchyma score of (**B**) 5 cm from the root tip and (**C**) 5 cm from the root base. The results represent the mean ± standard error of the mean of three independent biological replicates with three technical replicates each. Asterisk (*) indicates that the treatment was significantly different *(p* < 0.05).

The well-watered bananas recorded an average aerenchyma score of 1.17 to 1.75 throughout the experiment (Figures 2B and 2C). The aerenchyma size of the root tip did not show significant difference between well-watered and waterlogged samples (Figure 2B). However, the aerenchyma score of the root base of waterlogged samples was significantly higher than well-watered samples on days 14 and 24 (Figure 2C).

### 2.3. Proline and Malondialdehyde Contents Changed under Waterlogging Stress

Malondialdehyde (MDA) and proline contents were determined on banana root samples (Figure 3). MDA and proline contents remained relatively similar to well-watered samples throughout the first 5 days of the waterlogging treatment. However, significant changes in MDA content were observed after 7 days of waterlogging treatment (Figure 3).

**Figure 3.**
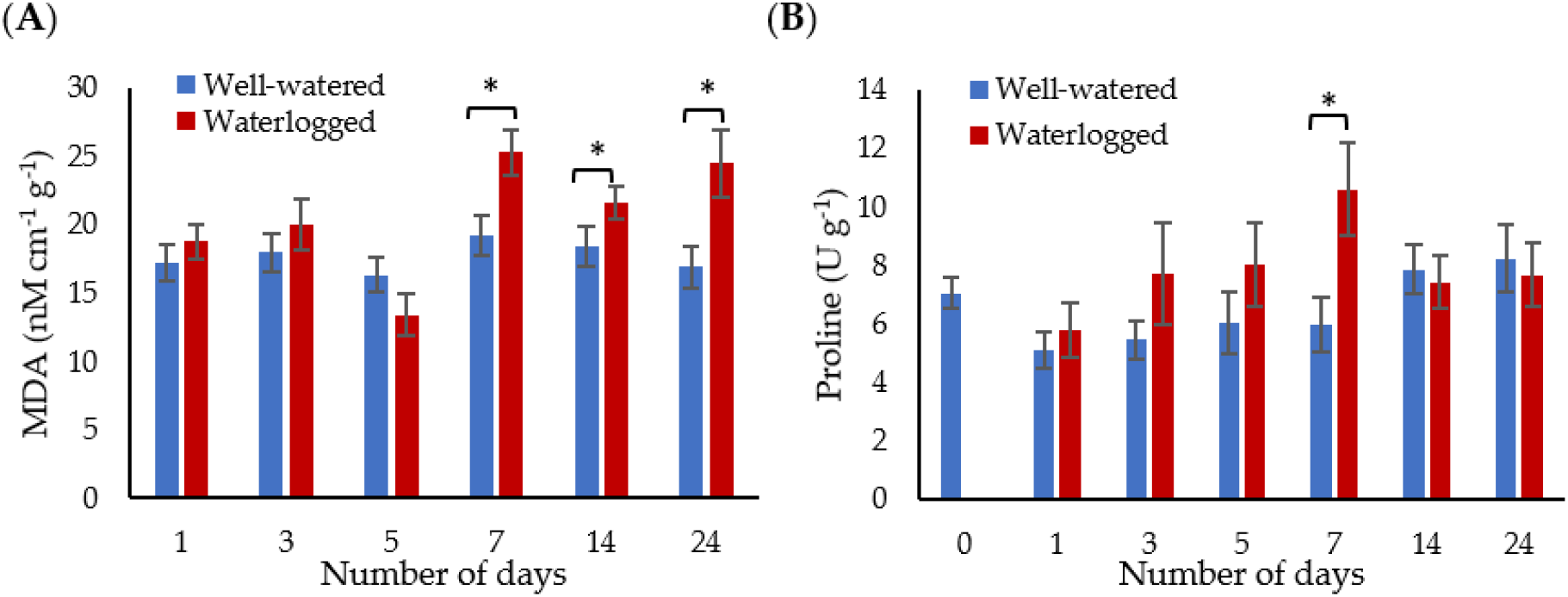
Osmolyte and lipid peroxidation changes in the well-watered and waterlogged banana plants. (**A**) Malondialdehyde (MDA) and (**B**) proline contents of banana roots. Error bars indicate the standard error of three independent replications. Asterisks (*) indicate a significant difference between well-watered and waterlogged samples at*p* < 0.05.

### 2.4. Waterlogging Stress Enhances Antioxidant Defense Systems

The antioxidant enzyme activities in well-watered and waterlogged banana plants were measured at 1, 3, 5, 7, 14, and 24 days to determine the ROS scavenging mechanism of bananas towards waterlogging stress (Figure 4). Waterlogging stress significantly increased the superoxide dismutase (SOD) activity, producing the highest level of 0.112 U g^-1^ after 1 day of waterlogging (Figure 4A). However, the SOD activity decreased to its lowest level (0.0136 U g^-1^) after 24 days of waterlogging. The ascorbate peroxidase (APX) level of the waterlogged samples was significantly higher than well-watered samples after 7, 14, and 24 days of waterlogging (Figure 4B). The glutathione reductase (GR) enzyme activity for well-watered samples increased from 0.148 U g^-1^ on day 1 to 0.676 U g^-1^ on day 24. In contrast, a significant increase in the GR enzyme activity for waterlogged samples was observed after 7 days, from 0.789 U g^-1^ to 1.045 U g^-1^ on day 24 (Figure 4C). The glutathione peroxidase (GPX) levels in waterlogged bananas were significantly higher than in well-watered plants on days 3, 7, and 24 of waterlogging, where the GPX level was highest on day 3 (0.287 μmol min^-1^ g^-1^) (Figure 4D). The catalase (CAT) activity was found to be similar for well-watered plants, with the highest recorded on day 7 for waterlogged bananas (0.120 nM cm^-1^ min^-1^ g^-1^) (Figure 4E). The H2O2 content of waterlogged samples was significantly higher than well-watered samples on day 14 and day 24 of waterlogging (Figure 4F).

**Figure 4.**
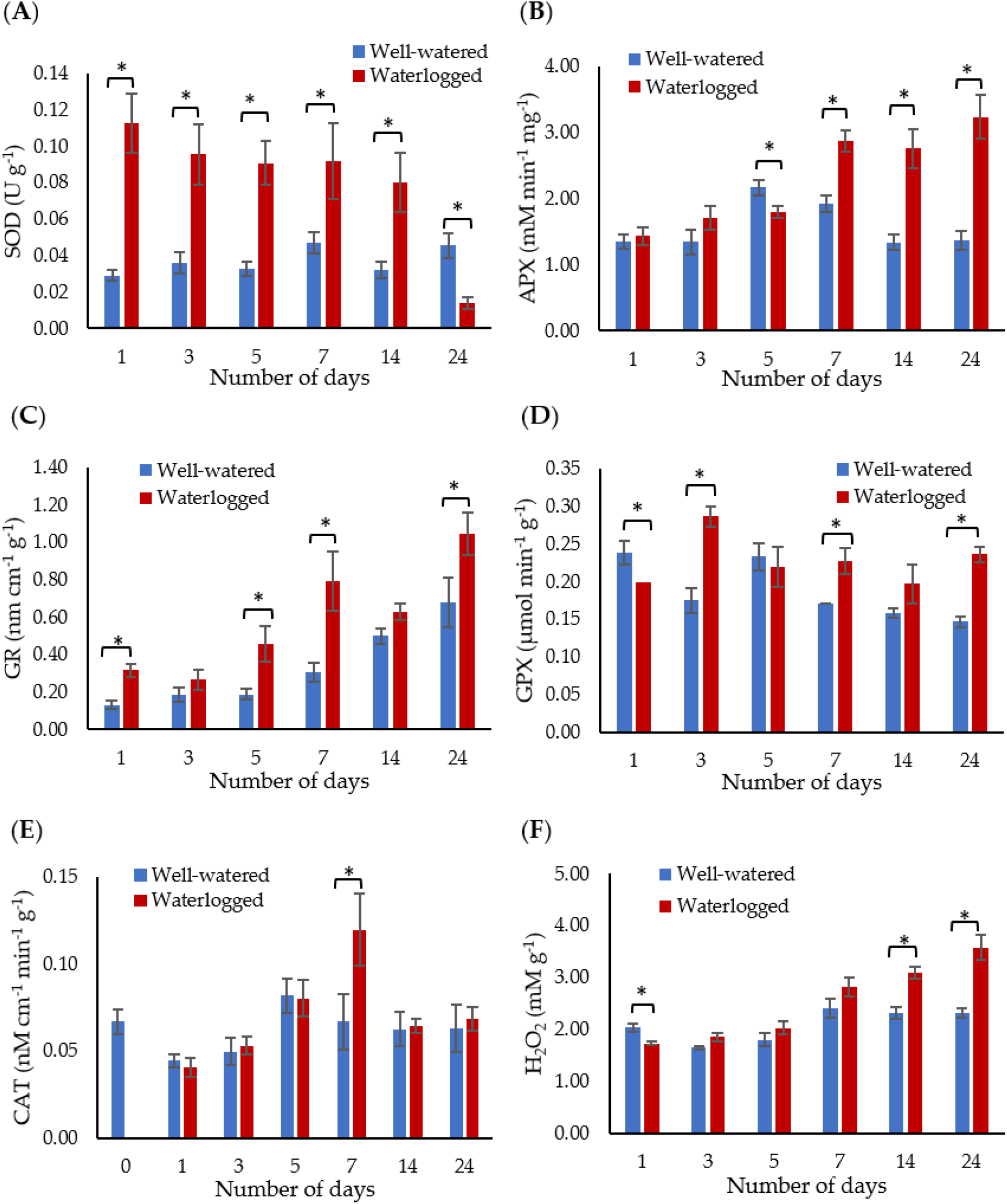
Activities of (**A**) superoxide dismutase (SOD), (**B**) ascorbate peroxidase (APX), (**C**) glutathione reductase (GR), (**D**) glutathione peroxidase (GPX), (**E**) catalase (CAT) and (**F**) hydrogen peroxidase (H2O2) in banana plants in response to waterlogging. Error bars indicate the standard error of three independent replications. Asterisks (*) indicate a significant difference between well-watered and waterlogged samples at *p* < 0.05.

### 2.5. Waterlogging Stress Altered Gene Transcription in Roots

We selected three *ERFVII,* namely *MaERFVII-1* (Macma4_09_g14450), *MaERFVII-2* (Macma4_08_g23860), and *MaERFVII-3* (Macma4_02_g02290), and *ADH1* (Macma4_02_g11450) genes for qPCR analysis to determine the early response of bananas toward waterlogging stress. Alcohol dehydrogenase (ADH) was selected as it plays a critical role in flooding tolerance. When banana plants were exposed to different waterlogging duration, we found that two out of three *ERFVII* genes were significantly *(p* < 0.05) upregulated on day 1 (Figure 5). The *ADH1* expression in waterlogged bananas was upregulated by 2.93-and 5.4-fold on day 1 and day 3, respectively, compared to well-watered bananas. We selected a 1-day waterlogging treatment for the subsequent RNA sequencing. The RNA samples with a RIN value of more than 7 were sequenced (Supplementary Table 2).

**Figure 5.**
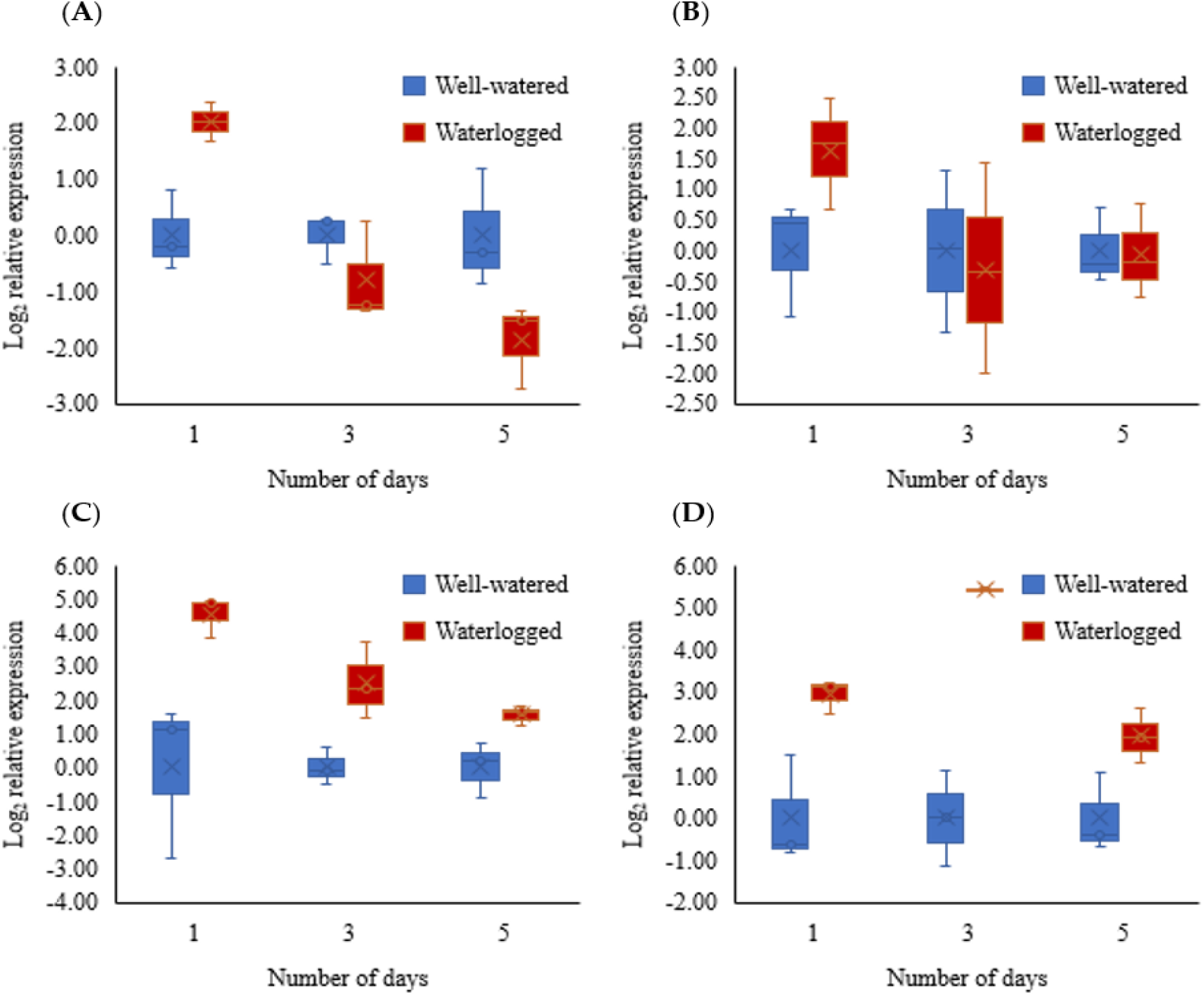
Gene expression of (**A**) *MaERFVII-1,* (**B**) *MaERFVII-2,* (**C**) *MaERFVII-3,* and (**D**) *ADH1* of banana for 1, 3, and 5 days of waterlogging. Error bars indicate standard error between biological replicates. Asterisks (*) indicate a significant difference between well-watered and waterlogged samples at *p* < 0.05.

The gene expression profiles of well-watered and waterlogged samples were analyzed using RNA-sequencing. Of the 407 million raw reads generated from the six cDNA libraries, approximately 406 million clean reads were obtained, ranging from 66.2 to 70.7 million reads per library (Supplementary Table 3). The clean reads were then mapped to the banana genome retrieved from the banana genome hub DH-Pahang (version 4.3) using Hisat2 (v2.0.1). The mapping rates of each library ranged from 80.37 % to 83.54 % (Supplementary Table 4). Of the 3,508 DEGs, a total of 1,470 were upregulated, and 2,038 were downregulated (Supplementary Table 5). The identified unigenes were classified according to their GO (Figure 6A). Most DEGs in the biological process category were involved in response to abscisic acid and salt stress and positive regulation of transcription. The cellular component category contained terms, such as ‘nucleus’, ‘extracellular region’, and ‘cell wall’. The most dominant terms in the molecular function category were ‘sequence-specific DNA binding transcription factor activity’, ‘transcription regulation’, and ‘heme binding’ (Table 1).

**Figure 6.**
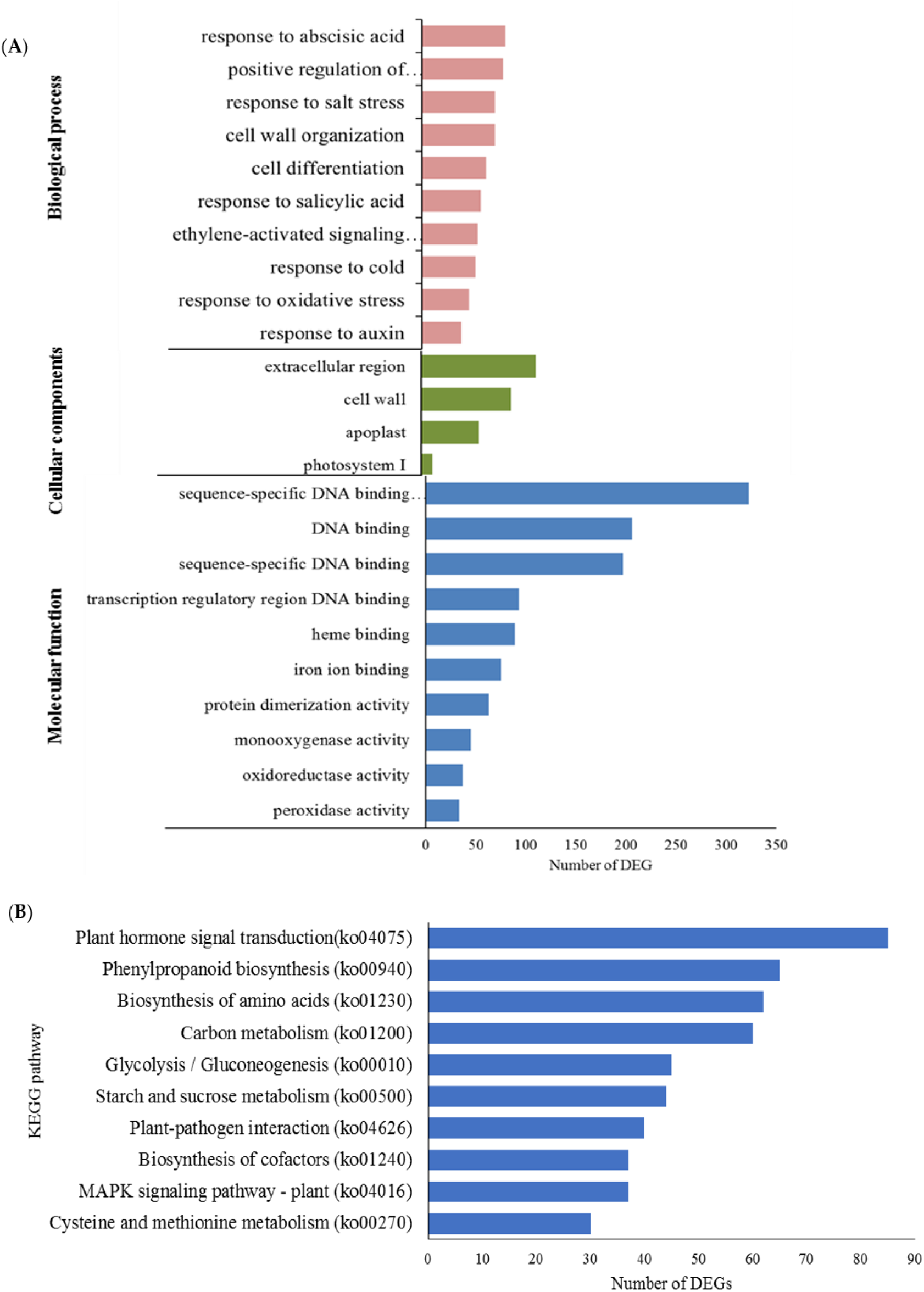
Functional categorization and enrichment of differentially changed genes between well-watered and waterlogged banana root samples. (**A**) Gene ontology enrichment according to biological processes, molecular functions, and cellular components and (**B**) the top ten significantly enriched KEGG pathways based on functional category.

There were 795 unigenes annotated according to the KEGG database [24], with 125 pathways significantly enriched. These include plant hormone signal transduction (ko04075), phenylpropanoid biosynthesis (ko00940), amino acid biosynthesis (ko01230), carbon metabolism (ko01200), and glycolysis (ko00010) (Figure 6B).

**Table 1.** The GO terms of banana 24 h after waterlogging stress.

| GO ID | GO term | p value | FDR | Upregulated DEG |
| --- | --- | --- | --- | --- |
| GO:0045893 | Positive regulation of transcription, DNA-templated | 5.40E-08 | 2.80E-05 | 38 |
| GO:0071555 | Cell wall organization | 3.37E-05 | 6.16E-03 | 38 |
| GO:0009737 | Response to abscisic acid | 7.20E-09 | 4.07E-06 | 31 |
| GO:0009873 | Ethylene-activated signaling pathway | 5.98E-08 | 2.86E-05 | 29 |
| GO:0009651 | Response to salt stress | 4.82E-04 | 4.83E-02 | 26 |
| GO:0006979 | Response to oxidative stress | 4.18E-06 | 1.24E-03 | 21 |
| GO:0009409 | Response to cold | 2.61E-04 | 3.12E-02 | 17 |
| GO:0010200 | Response to chitin | 4.45E-05 | 7.28E-03 | 16 |
| GO:0010411 | Xyloglucan metabolic process | 9.77E-06 | 2.43E-03 | 16 |
| GO:0042546 | Cell wall biogenesis | 4.29E-05 | 7.21E-03 | 16 |
| GO:0009751 | Response to salicylic acid | 1.17E-13 | 1.45E-10 | 15 |
| GO:0042744 | Hydrogen peroxide catabolic process | 1.22E-06 | 3.98E-04 | 14 |
| GO:0030154 | Cell differentiation | 9.60E-07 | 3.31E-04 | 12 |
| GO:0042545 | Cell wall modification | 2.81E-04 | 3.29E-02 | 12 |
| GO:0009611 | Response to wounding | 2.00E-06 | 6.22E-04 | 11 |
| GO:0055114 | Oxidation-reduction process | 4.77E-05 | 7.61E-03 | 11 |
| GO:0009753 | Response to jasmonic acid | 6.59E-06 | 1.86E-03 | 9 |
| GO:0009693 | Ethylene biosynthetic process | 2.65E-05 | 5.14E-03 | 9 |
| GO:0009835 | Fruit ripening | 1.59E-04 | 2.30E-02 | 9 |
| GO:0009741 | Response to brassinosteroid | 2.09E-04 | 2.74E-02 | 9 |
| GO:0009737 | Response to abscisic acid | 7.20E-09 | 4.07E-06 | 51 |
| GO:0030154 | Cell differentiation | 9.60E-07 | 3.31E-04 | 51 |
| GO:0009651 | Response to salt stress | 4.82E-04 | 4.83E-02 | 46 |
| GO:0009751 | Response to salicylic acid | 1.17E-13 | 1.45E-10 | 43 |
| GO:0045893 | Positive regulation of transcription,<br>DNA-templated | 5.40E-08 | 2.80E-05 | 42 |
| GO:0009409 | Response to cold | 2.61E-04 | 3.12E-02 | 36 |
| GO:0071555 | Cell wall organization | 3.37E-05 | 6.16E-03 | 34 |
| GO:0009733 | Response to auxin | 3.92E-04 | 4.06E-02 | 31 |
| GO:0009873 | Ethylene-activated signaling<br>pathway | 5.98E-08 | 2.86E-05 | 26 |
| GO:0006979 | Response to oxidative stress | 4.18E-06 | 1.24E-03 | 25 |
| GO:0009753 | Response to jasmonic acid | 6.59E-06 | 1.86E-03 | 25 |
| GO:0009611 | Response to wounding | 2.00E-06 | 6.22E-04 | 24 |
| GO:0009813 | Flavonoid biosynthetic process | 1.47E-05 | 3.20E-03 | 20 |
| GO:0055114 | Oxidation-reduction process | 4.77E-05 | 7.61E-03 | 18 |
| GO:0042744 | Hydrogen peroxide catabolic<br>process | 1.22E-06 | 3.98E-04 | 17 |
| GO:0071365 | Cellular response to auxin stimulus | 2.58E-07 | 1.07E-04 | 15 |
| GO:0031408 | Oxylipin biosynthetic process | 3.68E-05 | 6.53E-03 | 15 |
| GO:0010200 | Response to chitin | 4.45E-05 | 7.28E-03 | 14 |
| GO:1901332 | Negative regulation of lateral root<br>development | 2.33E-09 | 1.45E-06 | 12 |
| GO:0010345 | Suberin biosynthetic process | 1.34E-05 | 3.20E-03 | 11 |

### 2.6. Hormone Signaling Pathways are Differentially Regulated in Waterlogged Bananas

A total of 275 DEGs involved in plant hormone signaling transduction pathways were altered in waterlogged plants (Figure 7A). Specifically, 34.9 % were associated with ABA signaling, 24.6 % with ethylene signaling, 18.5 % with auxin signaling, 15.5 % with jasmonic acid, and 1.7 % with brassinosteroid (Figure 7A). Genes enriched in the ABA pathway include bZIP domain-containing protein, abscisic acid-insensitive 5-like protein, abscisic receptor PYL, and protein phosphatase 2C. Of the 96 genes involved in this pathway, 39 were upregulated, and 57 were downregulated (Supplementary Table 5).

**Figure 7.**
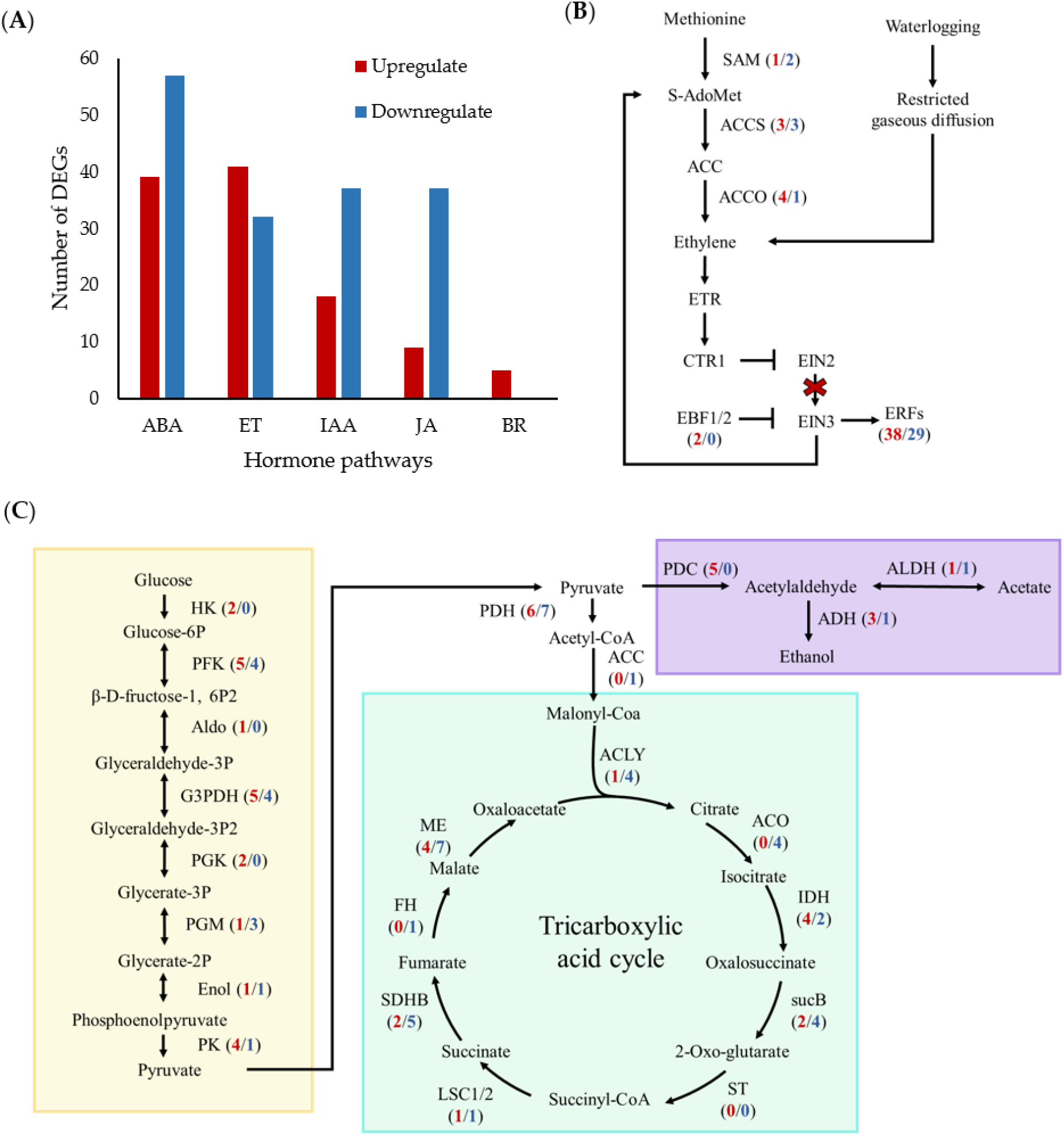
Plant hormone signal transduction pathway after 1 day of waterlogging stress. (**A**) The number of hormone-signal-related DEGs changes in bananas under waterlogging stress. The DEGs were involved in (**B**) the ethylene biosynthetic pathway and (**C)** carbon metabolism. The yellow box highlights the glycolysis pathway, leading to the ethanol fermentation in the purple box, and lastly, the blue box contains genes involved in the TCA cycle. The arrows indicate activation of the gene, while the blunt end represents inhibition of the gene. The red cross indicates the inactivity of the EIN3 pathway that leads to the activation of ERF TFs. The numbers in parentheses correspond to the number of upregulated (red) and down-regulated genes (blue). [ABA, abscisic acid; ACCO, 1-aminocyclopropane-1-carboxylic acid oxidase; ACCS, 1-aminocyclopropane-1-carboxylic acid synthase; ACLY, ATP citrate (pro-S)-lyase; ACO, aconitate hydratase; ADH, alcohol dehydrogenase; ALDH, aldehyde dehydrogenase; Aldo, aldolase; BR, brassinosteroid; CTR1, constitutive triple response 1; EBF, ethylene binding factor; ET, ethylene; EIN2/3, ethylene insensitive 2/3; Enol, enolase; IAA, auxin; FH, fumarate dehydrogenase; G3PDH, glyceraldehyde-3-phosphate dehydrogenase; HK, hexokinase; IDH, isocitrate dehydrogenase; JA, jasmonic acid; LSC1/2, succinate-CoA ligase; ME, malate dehydrogenase; PDC, pyruvate decarboxylase; PDH, pyruvate dehydrogenase; PEPC, phosphoenolpyruvate carboxykinase; PFK, phosphofructokinase; PGK, phosphoglycerate kinase; PGM, phosphoglycerate mutase; PK, pyruvate kinase; SAM, S-adenosyl-L-methionine; SDHB, succinate dehydrogenase; ST, succinyl transferase; sucB, 2-oxoglutarate dehydrogenase].

In the ethylene signaling pathway, upregulated genes include ethylene biosynthetic genes (*ACCO* and *ACCS*), ethylene transport receptors, and ethylene-responsive transcription factors (TFs) (Figure 7B). Enrichment analysis revealed that auxin-responsive genes, such as *SAUR32* (Macma4_02_g06820, Macma4_07_g20980), *IAA30,* and *IAA4,* were downregulated. However, the auxin biosynthesis gene, namely *IAA-amino synthetase GH3.8* (Macma4_02_g16920 and Macma4_05_g01480), was upregulated by 2.4-and 3.4-log2 fold change. These results indicated that waterlogging could lead to a different hormone signaling regulation, which might play an important role in the waterlogging response and adaptation in bananas.

### 2.7. Hypoxia-Related Genes Showed Transcriptional Responses in Waterlogged Bananas

To understand the changes in waterlogging response in banana plants, pathways related to hypoxia and carbon metabolism were assessed. All eight genes related to oxygen sensing, the plant cysteine oxidase (PCO) gene, were significantly upregulated by 1.45-to 5.0-fold in waterlogged samples (Supplementary Table 5).

For carbohydrate metabolism, most genes related to glycolysis, including hexokinase, phosphofructokinase, aldolase, enolase, and glyceraldehyde-3-phosphate dehydrogenase, were upregulated (Figure 7C). The genes related to ethanol fermentation were upregulated in response to waterlogging, i.e., five pyruvate decarboxylase (PDC) genes and three *ADH* genes. In contrast, most tricarboxylic acid (TCA) cycle genes, namely citric synthase, aconitate hydratase, 2-oxoglutarate dehydrogenase, succinate dehydrogenase, fumarate dehydrogenase, and malate dehydrogenase, were downregulated. These results indicate that fermentation is critical for bananas to respond to early waterlogging stress.

### 2.8. Expression of Transcription Factors (TF) Affected in Waterlogged Bananas

Among 3,508 DEGs, a total of 267 were TFs, with 113 TFs being upregulated and 154 TFs being downregulated *(p* adjust < 0.01) (Figure 8). These TF families include AP2/ERF, bHLH, MYB, NAC, bZIP, and WRKY. Among ERF TFs, 38 were upregulated, whereas 29 TFs were downregulated (Figure 8). ERFVII gene (Macma4_11_g05400) was the most upregulated gene with a log2 fold change of 4.4, indicating that this TF might be critical in waterlogging responses (Supplementary Table 5).

**Figure 8.**
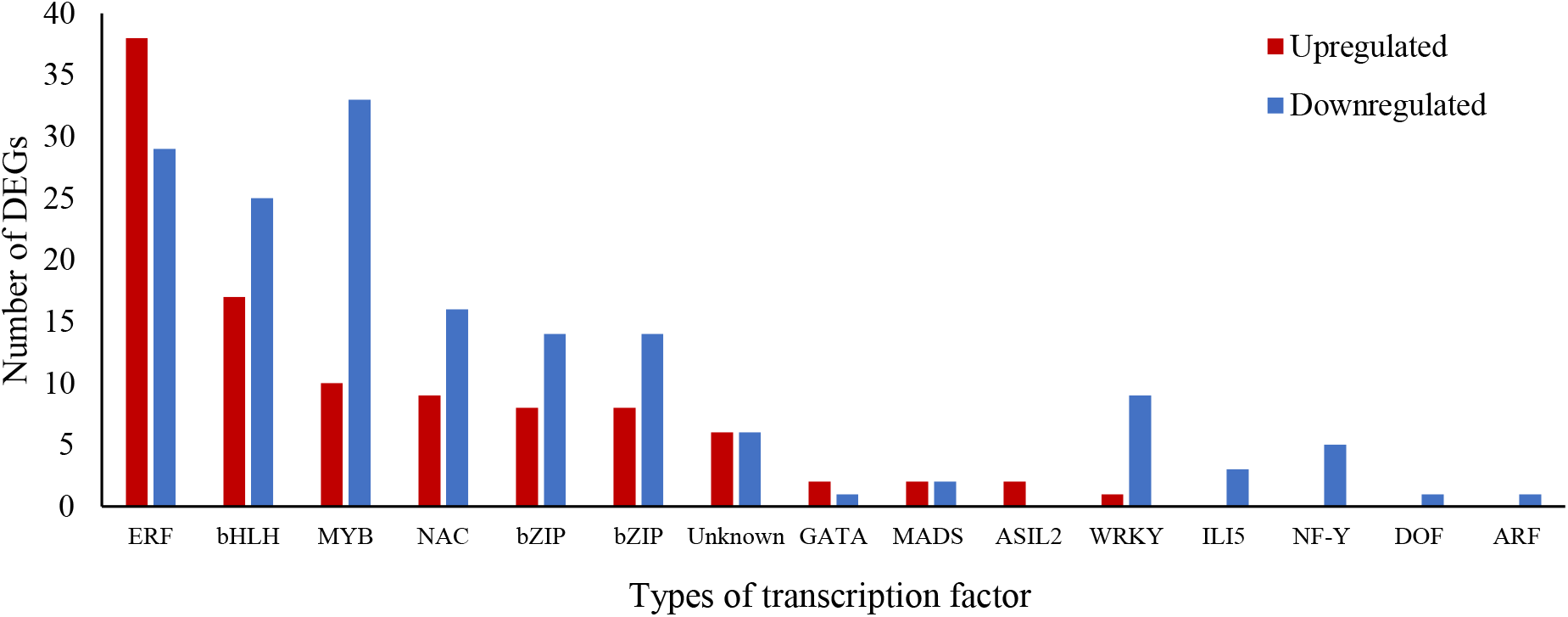
The number of waterlogging-related transcription factors that are classified according to their protein families. [ARF, auxin response factor; bHLH, basic helix-loop-helix; bZIP, basic zinc leucine zipper domain; DREB, dehydration-responsive element binding; ERF, ethylene response factor; MYB, myeloblastosis; NAC-NAM, ATAF, and CUC; ZF-TF, zinc finger transcription factor].

### 2.9. qPCR Validation of DEGs from the RNA-Sequencing

To validate the transcriptomic data, nine genes involved in the abscisic acid (ABA) biosynthetic pathway, ethylene signaling pathway, and a hypoxia-responsive gene were selected for qPCR analysis (Figure 9A). The expression of the selected genes between RNA-sequencing and qPCR were highly correlated, with a correlation coefficient of 0.9408 (Figure 9B).

**Figure 9.**
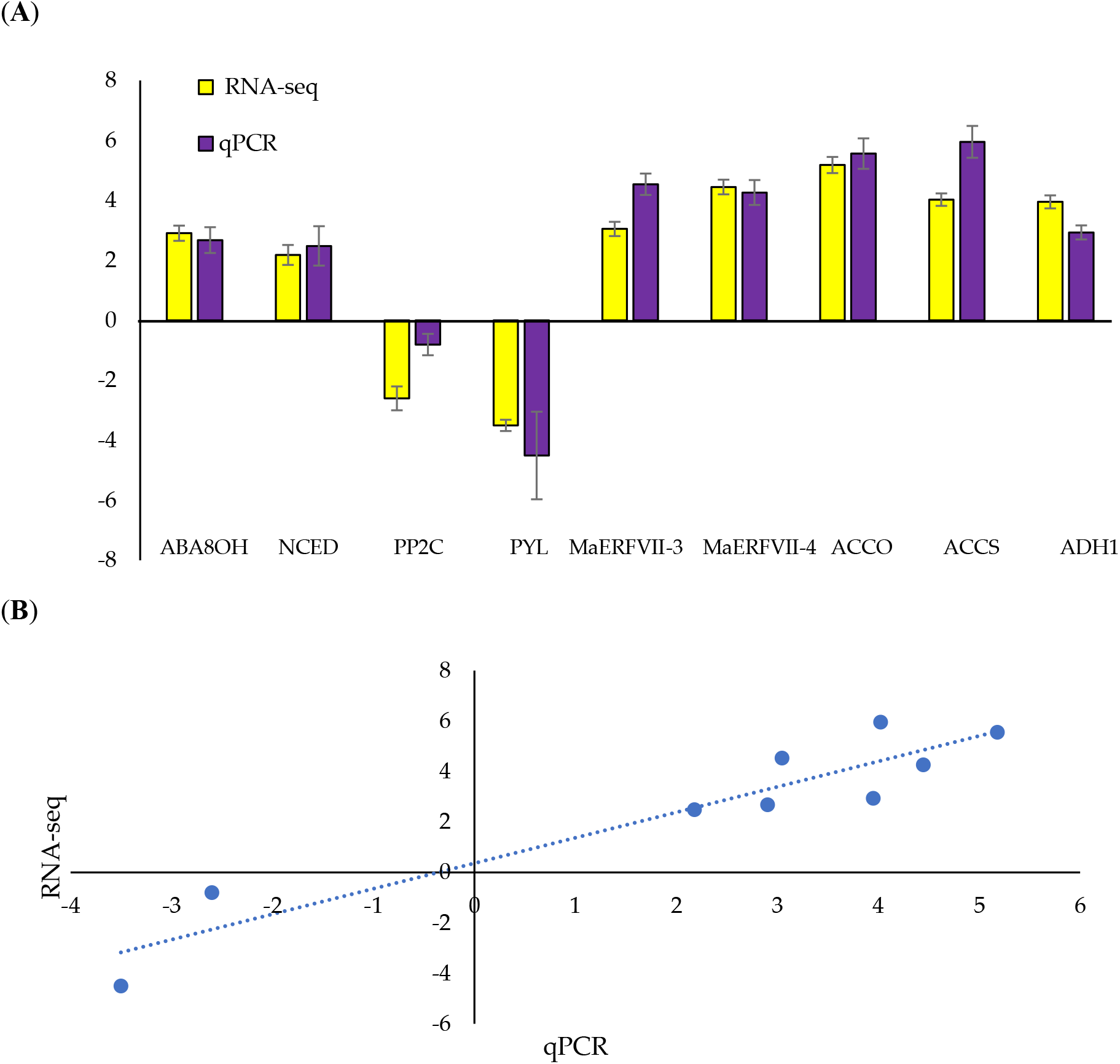
Comparison of RNA-sequencing and qPCR analyses. (**A**) The qPCR validation of up-and down-regulated differentially expressed genes in waterlogged bananas relative to the well-watered bananas. Values are means ± SE of three biological replicates in qPCR. The expression levels of each gene are expressed relative to the mean values of the control samples. (**B**) The correlation between qPCR and RNA-sequencing data. [ABA8OH, abscisic acid 8’ hydroxylase; ACCO, 1-aminocyclopropane-1-carboxylic acid oxidase; ACCS, 1-aminocyclopropane-1-carboxylic acid synthase; ADH1, alcohol dehydrogenase 1; NCED, 9-cis-epoxycarotenoid dioxygenase; PYL, PYR-like proteins; PP2C, type 2C protein phosphatase].

## 3. Discussion

### 3.1. Morphological Responses of Bananas Subjected to Waterlogging

As sessile organisms, plants are unable to escape when challenged by unfavorable environmental conditions. However, they can deploy complex morpho-physiological and molecular mechanisms to cope and adapt to these adverse environmental stresses [11]. In this study, the banana plants elicited morphological responses to combat the waterlogging stress. When exposed to excessive water, banana plants change their morphology to increase their chances of survival. Waterlogged soils progressively exert a detrimental effect on the plant root respiration and nutrient uptake. This phenomenon resulted in leaf wilting, which was observed at the latter stages of waterlogging. Although waterlogging did not affect the leaf area and RWC, it induced adventitious roots and aerenchyma formation. The results concurred with several previous studies that showed that waterlogging significantly increases the number of adventitious roots and aerenchyma formation [25,26]. Adventitious roots and aerenchyma are critical to improving the waterlogging tolerance in plants [27]. Waterlogged soils cause ethylene entrapment and affect auxin transportation, triggering the formation of adventitious roots [28]. The formation of aerenchyma is essential in plant waterlogging tolerance as it can provide oxygen to the submerged organs of a plant.

### 3.2. Antioxidant Changes in Roots of Waterlogged Bananas

Waterlogging stress generally leads to the accumulation of MDA and H2O2. Our findings agree with the previous studies, where MDA and H2O2 were elevated under waterlogging or hypoxia stress in various plants [29,30]. The overproduction of ROS induces lipid peroxidation. MDA is the end product of lipid peroxidation that has been frequently used to determine ROS-induced oxidative injury [31]. In this study, the significant differences in MDA levels at various treatment times indicate that MDA levels in bananas are time-specific. Proline is a ROS scavenger because it may help maintain protein structural stability [32], and waterlogging stress induces high proline accumulation in various plants, such as peach and cucumber [33,34].

The excessive accumulation of ROS accompanied by increasing antioxidant enzyme activities. In the present study, the increased antioxidant enzyme activities in the banana roots under waterlogging indicated that they possess efficient ROS scavenging systems. For example, the SOD activity remained high since day 1 of waterlogging, reaffirming the role of SOD as the first line of defense against ROS. In plants, APX and CAT are predominant enzymes in alleviating the damage caused by stresses [29]. This study recorded a remarkable increase in APX activity in waterlogged banana roots. Similar findings were also reported by [35] and [36], where the waterlogging stress increased APX activity. In contrast, the CAT activity for well-watered and waterlogged samples was not significantly different. GPX is an oxygen radical scavenger, catalyzing the reduction of H2O2 via glutathione to generate water and glutathione disulphide. Consequently, GR reduces glutathione disulphide to glutathione, supplying electron donors for the subsequent detoxification of H2O2 [37]. Our results showed that GPX and GR activities in waterlogged banana roots were generally higher than in well-watered roots, which are supported by other studies [38,39]. Taken together, the enhanced antioxidant enzyme system helps banana plants cope with accumulated ROS under waterlogging conditions.

### 3.3. Oxygen Sensing of Banana under Waterlogging Stress

Waterlogging affects oxygen-sensing in plants. PCO is an oxygen sensor in the N-degron pathway. Under normal conditions, ERFVII TF N-terminal cysteine residue is subjected to O2-dependent degradation through the N-end rule pathway by PCO [40]. In the absence of oxygen, PCO cannot destabilize ERFVII TFs, leading to the activation of hypoxia-responsive genes [41]. Hence, PCOs and ERFVIIs represent the oxygen-sensing machinery for regulating flooding survival in plants. However, in this study, six PCO genes in banana roots were upregulated under waterlogging stress. This might be due to the residual oxygen molecules present in the plants or water. Loreti et al. [42] showed that the *PCO2* expression in *A. thaliana* was upregulated under anoxia stress but reduced to a minimum level after 24 h of treatment.

ERFs are a critical flooding-responsive mechanism in plants. They are constitutively expressed but are degraded in the presence of oxygen [43]. Our results showed that among 67 ERF TFs, those in the IIa, IIIb, Va, VII, VIIIa, IX, and X subgroups were upregulated, while ERF TFs in IIb, Vb IXc, and Xa subgroups were downregulated. Both IIb and IIId subgroups contained upregulated and downregulated transcripts. From our transcriptomics results, six out of 18 genes of the ERFVII family were upregulated. Members of the ERFVII are well-known as the main regulator of plant responses to low oxygen conditions [44]. For instance, knockout of *erfVII* in *A. thaliana* decreased its tolerance to hypoxia [45]. In wheat, the overexpression of ERFVII improved the waterlogging tolerance without grain yield penalty [46].

It has been well documented that ethylene accumulation is a key signal for flooding. Our transcriptome data show several ethylene biosynthesis genes, such as S-adenosyl-L-methionine (SAM) and 1-aminocyclopropane-1-carboxylic acid oxidase (ACCO), were upregulated in the submerged roots. These results agree with the previous studies, showing that waterlogging stress promotes ethylene biosynthesis [47]. However, although ethylene biosynthesis genes were upregulated, the downstream genes in the ethylene pathway, such as *ETR, CTR1,* and *EIN3,* were not affected by waterlogging stress. The increased EBF transcription led to EIN3 inhibition. Nevertheless, this inhibition did not decrease the transcription of ERF TFs. There is growing evidence of the EIN3/EIL1-independent mediated pathway in the ethylene pathway. For example, the double knockout *ein3ein1* Arabidopsis mutants were able to respond to ethylene under stress [48]. Hence, we speculate that an EIN3-independent pathway might be involved in activating ERF TF transcription. However, further studies are required to verify this.

### 3.4. ABA Pathways of Bananas under Waterlogging Stress

In waterlogged banana roots, many genes associated with the ABA pathway were found to have a significant transcriptional response to the waterlogging stress. ABA is a phytohormone known to respond to drought, salinity, and cold stress. The reduction in ABA content is a common response to waterlogging. In our study, the gene encoding the rate-limiting enzyme in ABA biosynthesis, 9-cis-epoxycarotenoid dioxygenase 1 (NCED1), was downregulated, while the ABA degradation enzyme, ABA 8’-hydroxylase gene, was upregulated. Several studies have demonstrated that ABA is involved in regulating adventitious root and aerenchyma development in submerged plants [49, 50]. ABA content decreased following the increase in ethylene concentration under waterlogging. For example, the exogenous ABA treatment inhibited adventitious root primordia formation in tomatoes [49] and aerenchyma formation in soybean roots [51]. A decrease in ABA is essential for waterlogged-induced gibberellic acid to promote shoot elongation [52]. These results indicate that ABA acts as a negative regulator in waterlogging tolerance in plants.

### 3.5. Carbon Metabolism is Essential for Energy Supply during Waterlogging

When plants are exposed to hypoxic conditions, they immediately alter their gene transcription to synthesize anaerobic polypeptides, derived from glycolysis and ethanol fermentation [53]. This study found that glycolysis-related genes were significantly expressed on 1 day of waterlogging. These include hexokinase, phosphoglycerate kinase, and pyruvate kinase. Nutrient uptake under waterlogging conditions is inhibited for hypoxic roots. Given the need to maintain sufficient energy for plant survival, it is not surprising that carbon metabolism was differentially altered under waterlogging stress. Previous studies showed that waterlogging stress increases starch metabolism and the glycolysis/gluconeogenesis pathways in the waterlogged roots of *Cerasus sachalinensis* [54] and barley [55].

Aerobic respiration in roots is limited under low oxygen conditions, resulting in glycolysis being channeled to fermentative pathways. These changes are necessary to produce energy for cell survival. In roots, the expression of key genes involved in alcohol fermentation was upregulated under waterlogging conditions. Indeed, PDC and ADH were upregulated in waterlogged bananas. Pyruvate generated from glycolysis is metabolized to ethanol via PDC and ADH. This process releases energy in the form of adenosine triphosphate (ATP) to increase the chances of plant survival in the absence of aerobic respiration.

The changes in carbon metabolism are often accompanied by the changes in nitrogen metabolism to maintain the carbon-nitrogen balance. Genes associated with the TCA were downregulated in the roots of waterlogged banana plants. Pathways that provide intermediates for the TCA cycle, such as glutamate synthase (GS), glutamate dehydrogenase (GDH), and glutamate carboxylase, were downregulated under waterlogging stress. Similarly, waterlogging stress reduced the activity of GS and GDH in the stem and leaves of maize [56]. The increase of amino acid catabolism is required to compensate for the insufficient energy supply during stress. This can be seen in the waterlogged banana roots, where the alanine biosynthesis gene (alanine aminotransferase; AlaAT) was upregulated. Alanine is a carbon and nitrogen source, and its biosynthesis is crucial for plants to survive waterlogging [57]. These results suggest that activation of amino acid catabolism and alcohol fermentation are vital to fulfilling the energy demand in plants under waterlogging stress.

### 3.6. A Model for the Response of Banana Plants to Waterlogging

We propose a model for waterlogging responses in banana plants (Figure 10). Waterlogging affects changes in plant cells, such as limitation of gas diffusion, nutrient depletion, and increased ROS production. To respond and adapt to waterlogging, banana plants have evolved to rapidly elongate their adventitious roots and aerenchyma formation and activate the antioxidant defense system. Our data revealed transcriptional changes in the downstream components of hormone signaling and carbon and energy metabolisms during waterlogging.

**Figure 10.**
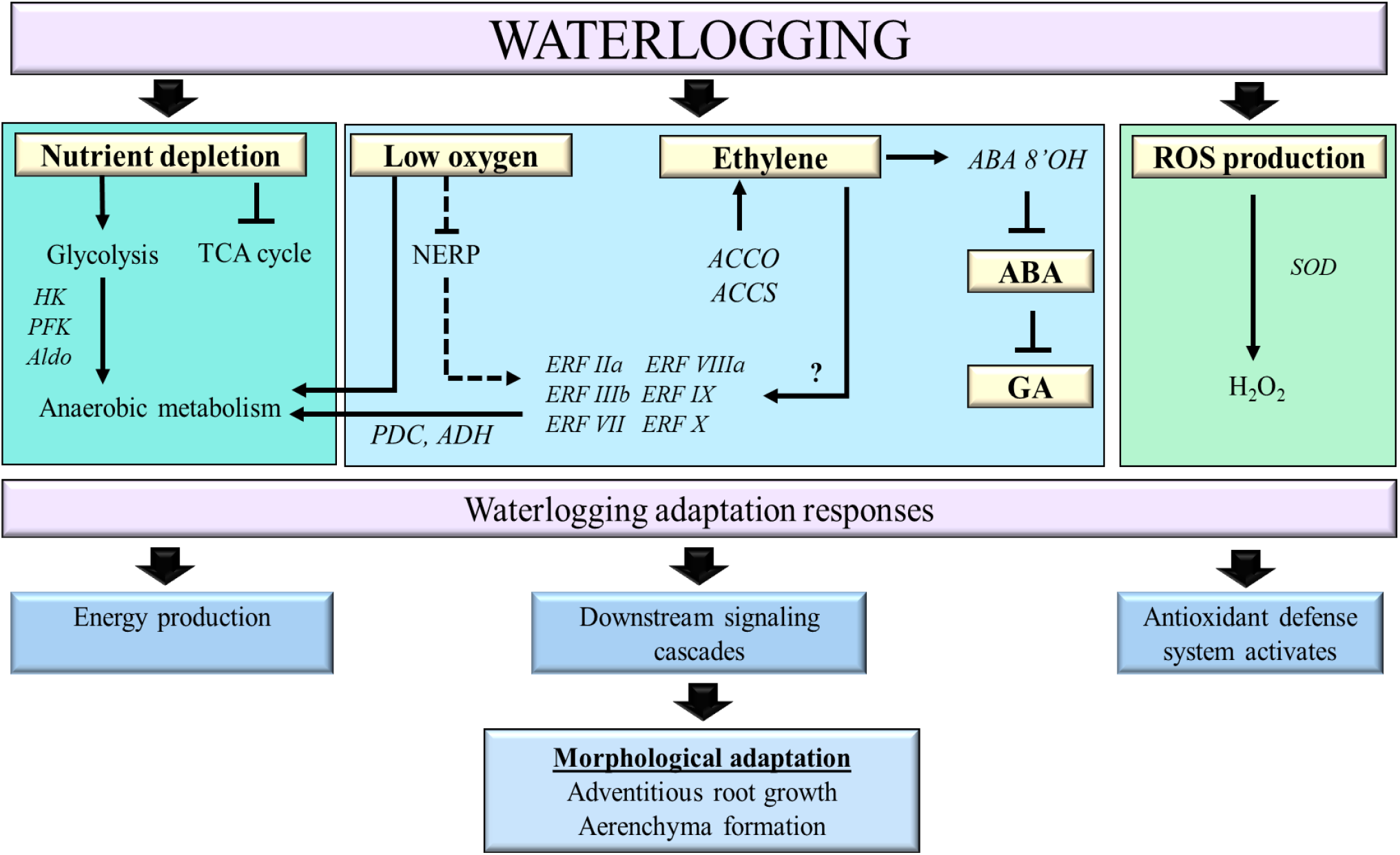
The proposed model of the pathways involved in *M. acuminata* under waterlogging stress. Arrows indicate the activation of the pathway, while blunt end arrows indicate inhibition of the pathway. The dashed line indicates partial activation of the NERP pathway, leading to the partial inhibition of ERFVII TFs. The question mark indicates an unknown EIN3/EIL independent pathway that activates ERF genes. [ABA, abscisic acid; ABA 8’OH, abscisic acid 8’ hydroxylase; ACCO, 1-aminocyclopropane-1-carboxylic acid oxidase; ACCS, 1-aminocyclopropane-1-carboxylic acid synthase; Aldo, aldolase; ERF, ethylene response factor; GA, gibberellic acid; H2O2, hydrogen peroxide; HK, hexokinase; NERP, N-end rule pathway; PDC, pyruvate decarboxylase; PFK, phosphofructokinase; SOD, superoxide dismutase; TCA, tricarboxylic acid].

## 4. Materials and Methods

### 4.1. Plant Material and Waterlogging Treatment

Two-month-old banana (*M. acuminata* cv. Berangan) seedlings purchased from the MB Tissue Culture Resources, Selangor, were acclimatized in a greenhouse at the Universiti Malaya. The plants were maintained under natural light (12 h photoperiod) at a controlled temperature (25 ± 2 ^o^C) under a relative humidity of 65 ± 5 %. Waterlogging treatment was conducted by filling the plastic containers with water 2 cm above the soil surface. Each plastic container contained five banana plants. The containers were refilled with tap water to maintain a constant water level. The well-watered bananas were planted in well-drained soil and watered once a day during treatment. The waterlogging treatments were conducted for 1, 3, 5, 7, 14, and 24 days. Each time point consisted of nine plants (n = 9). The morphological changes of the treated and non-treated banana plants were determined for each time point. After waterlogging treatment, root samples were harvested, ground into a fine powder in the presence of liquid nitrogen, and stored at-80 ^o^C until use. These samples were used for the subsequent gene expression analysis and biochemical assays.

### 4.2. Morphological Analysis

Morphological changes, such as total root length, R/S DW ratio, leaf area, and leaf RWC, were measured. The total root length was determined based on the sum of the length of each root. Each root was measured using a meter ruler. The plants were divided into the shoot and root parts and oven-dried at 80 ^o^C for 2 weeks or until constant weight for dry mass. The R/S DW ratio was calculated based on the dry matter of the root and shoot parts of the plants.

The leaf area differences were calculated based on the differences between leaf area measurements before and after the waterlogging treatment. The RWC was measured according to Turner [58].For RWC, the first mature leaf was excised into 2 cm ×2 cm size before measuring the total fresh weight. Next, leaves were submerged in distilled water, and saturated weight was measured after 24 h. Leaves were dried in a 60 ^o^C oven for 24 h before measuring DW. RWC was calculated according to the following equation:

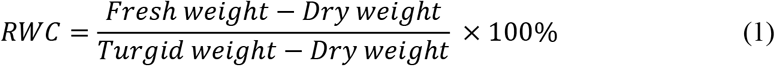

### 4.3. Adventitious Root and Aerenchyma Formation

Adventitious roots were calculated based on the number of roots per plant protruding from the banana stem. Aerenchyma formation was assessed according to Mano et al. [59]. A total of three lateral roots with a root length of more than 15 cm were randomly selected from each banana plant in each treatment group to determine the presence of aerenchyma. Cross-sections of roots about 1 cm thick were made using a handheld razor blade 5 cm beyond the root base and 5 cm beyond the root tip (Figure 11A). Aerenchyma in the root cortex was visually scored under an inverted fluorescent microscope (Olympus, Japan) on a scale of 0 to 3, where 0 indicates no aerenchyma, 1 indicates radial aerenchyma formation, 2 indicates radial formation extended to the epidermis, and 3 indicates well-formed aerenchyma (Figure 11B).

**Figure 11.**
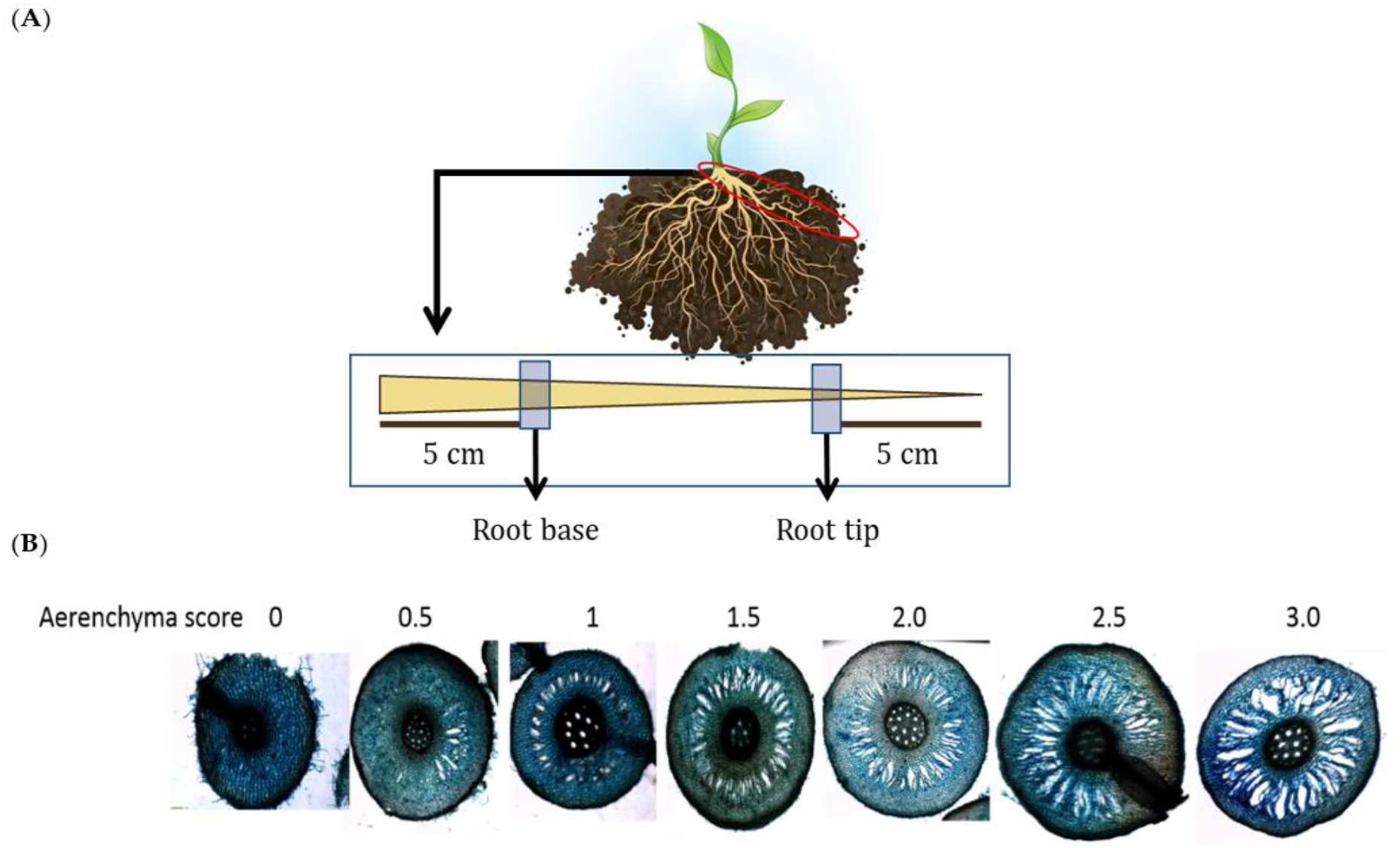
Determination of root aerenchyma formation. (**A**) The roots were removed from the plant and excised 5 cm from the root base, which is closer to the stem, and 5 cm at the root tip, which is away from the plant. (**B**) The aerenchyma score was assessed according to the descriptions from Mano et al. [59].

### 4.4. Malondialdehyde, Proline, and Hydrogen Peroxidase

H2O2 content was measured as described by Junglee et al. [60]. Frozen banana root tissues (150 mg) were homogenized with 1 ml of buffer solution containing 0.025 % (w/v) TCA, 0.5 M potassium iodide, and 2.5 mM potassium phosphate buffer. The absorbance of the supernatant was measured at 350 nm using a spectrophotometer.

Malondialdehyde (MDA) content was determined as described by Amnan et al. [31].The root tissues were homogenized and incubated with 0.5 % (w/v) thiobarbituric acid (TBA) in 20 % (w/v) trichloroacetic acid (TCA) at 95 ^o^C for 15 min. The solution was then placed on ice to terminate the reaction and centrifuged to collect the supernatant. A spectrophotometer was used to measure the absorbance of the solution at 532 and 600 nm, where the net absorbance value was recorded. Lastly, the MDA content was calculated with the extinction-coefficient 155 mM^-1^ cm^-1^.

Proline content was estimated as described by Bates et al. [61]. A total of 0.5 g samples were homogenized in 1 ml of 70 % ethanol. The filtrate was mixed with an equal volume of acid-ninhydrin and acetic acid for 1 h at 100 ^o^C. The reaction was terminated using an ice bath. The solution was partitioned against 4 ml toluene and measured in the organic layer for absorbance at 520 nm. Commercially available proline was used as a standard to construct a calibration curve.

### 4.5. Antioxidant Enzyme Assays

The antioxidant enzyme activities, namely SOD, APX, GR, GPX, and CAT, in well-watered and waterlogged-banana plants, were measured. The antioxidant enzyme activities were determined according to Zhang et al. [62].In brief, soluble proteins were extracted from the homogenized roots in potassium phosphate buffer (pH 7.0) containing ethylenediaminetetraacetic acid (EDTA) and polyvinylpyrrolidone, with the addition of ascorbic acid in the case of APX assay. The supernatant was collected through centrifugation and used for SOD, APX, CAT, GR, and GPX assays.

Total SOD activity was monitored through the inhibition of photochemical reduction of nitro blue tetrazolium. The reaction mixture was illuminated for 10 min under a 35 W fluorescent tube. The reaction was then terminated by placing the tube in a dark container. The absorbance of the mixture at 560 nm was measured for all samples at 15 s intervals for 3 min.

APX activity was determined from the decrease in absorbance at 290 nm reading for 1 min in a reaction mixture. GR assay was performed according to Carlberg et al. [63] through the oxidation of 1 μmol of NADPH in 3 min at 290 nm. GPX was measured as the oxidation of guaiacol in the presence of H2O2. The reaction started after adding guaiacol followed by the changes of 470 nm over 3 min [64]. CAT activity was monitored through the consumption of H_2_O_2_ with an extinct coefficient of 39.4 mM^-1^ cm^-1^, resulting in a decrease in absorbance at 240 nm for 3 min.

### 4.6. RNA Extraction

Total RNA was extracted from the roots using the CTAB extraction method according to Asif et al. [65].The RNA quality was analyzed using Nanophotometer Perl^®^ (Implen, Germany), whereas the RNA integrity was visualized using agarose gel electrophoresis and Bioanalyzer. The extracted RNA was treated with DNase (Qiagen, Germany) according to the manufacturer’s protocol to remove residual genomic DNA.

### 4.7. Quantitative Real-Time PCR

Quantitative real-time PCR (qPCR) was performed to analyze the expression of ERFVII TFs. The first strand of cDNA was synthesized from total RNA (2 μg) using the NxGen M-MuLV Reverse Transcriptase cDNA synthesis kit (Lucigen, UK). The qPCR consisted of a final volume of 15 μl containing 10 ng of cDNA, 0.3 μM primers, and 2 × SG Fast qPCR Master Mix (Shanghai, China), with the elongation factor 1-alpha and tubulin as reference genes (Supplementary Table 1). The qPCR was carried out according to the manufacturer’s protocol. Relative expression levels were calculated according to Pfaffl[66].The qPCR analysis was carried out with three biological replicates and three technical replicates for each gene.

### 4.8. RNA Sequencing

The extracted RNA with a RIN value between 6.5 and 10 was used for poly-A enriched library construction. RNA sequencing was performed on an Illumina system with paired-end 150 bp reads. Raw reads were assessed for their quality using the FastQC program [67] and filtered using Cutadapt (v1.9.1) [68]. Cutadapt software was used to remove primer and adapter sequences from the raw reads to obtain high-quality clean data. The clean reads of all samples were aligned against the banana reference genome *(M. acuminata* DH-Pahang version 4.3) (https://banana-genome-hub.southgreen.fr/content/download) [69]. The mapping was performed using Hisat2 (v2.0.1) [70] using the default parameters. Differentially expressed genes (DEGs) were identified using the DESeq2 Bioconductor package [71]. GOSeq (v1.34.1) [72] was used to identify Gene Ontology (GO) terms that annotate a list of enriched genes, while the Kyoto Encyclopedia of Genes and Genome (KEGG) [24] was used to enrich genes in the KEGG pathway. The generated FASTQ files were deposited at the NCBI Sequence Read Archives database (https://www.ncbi.nlm.nih.gov/sra) under BioProject ID: PRJN850880.

### 4.9. Statistical Analysis

Statistical analysis was conducted with SPSS 23.0 (SPSS, Chicago, USA). All data were statistically analyzed by two-way analysis of variance (ANOVA) followed by Duncan’s multiple range test at a significance level of *p* <0.05.

## 5. Conclusions

This study describes the morphological and molecular responses of banana plants in response to waterlogging stress. In general, the waterlogging-stressed plants showed a higher number of adventitious roots, enlarged root aerenchyma, and enhanced antioxidant enzyme activities than well-watered plants. The early waterlogging-responsive mechanisms of bananas were determined by examining their transcriptome changes. Several unigenes were validated to be differentially expressed, and the majority of them were classified as abiotic stress-related transcription factors, signal transduction, and carbohydrate metabolisms. Although the ethylene biosynthetic pathway was upregulated, the ethylene-independent pathway activating ERF TFs might also be involved. Our findings provide insight into the complex molecular events involved in response to waterlogging stress of banana roots, which could help develop waterlogging resilient crops for the future climate.

## Supplementary Materials

Figure S1: Representative image of banana plants exposed to waterlogging stress for 0, 1, 3, 5, 7, 14, and 24 days.

Supplementary Table S1: Sequences of primers, banana genome hub gene ID, and the product length of the targets.

Supplementary Table 2: The RIN value, 28S/18S ratio and concentration of RNA prior library preparation. Supplementary Table S3: Summary of the raw and cleaned reads.

Supplementary Table S4: Total reads and mapped reads for well-watered and waterlogged bananas.

Supplementary Table S5: List of significant differentially expressed genes. Genes related to the transcription factor family and hormonal pathways were included in the table.

## Author Contributions

Conceptualization, B.C.T; methodology, E.Y.T., T.L.P. B.C.T; formal analysis, E.Y.T.; writing—original draft preparation, E.Y.T, B.C.T; writing—review and editing, B.C.T, C.H.T, B.N.A.; supervision, B.C.T, C.H.T, B.N.A., T.L.P.; funding acquisition, B.C.T. All authors have read and agreed to the published version of the manuscript.

## Funding

This research was funded by the Fundamental Research Grant Scheme (FP065-2018A), Royal Society-Newton Advanced Fellowship (IF004-2018), and Universiti Malaya RU Fund (ST003-2021).

## Data Availability Statement

Sequencing data are available at the National Center for Biotechnology Information (NCBI) under BioProject ID: PRJNA850880.

## Acknowledgments

Not applicable.

## Conflicts of Interest

The authors declare no conflict of interest.

## Notes

https://www.ncbi.nlm.nih.gov/bioproject/

### Competing Interest Statement

The authors have declared no competing interest.

## References

1. FAOUN. The future of food and agriculture: Trends and challenges. 2017.

2. Wang, X.; Xia, J.; Dong, B.; Zhou, M.; Deng, S. Spatiotemporal distribution of flood disasters in Asia and influencing factors in 1980–2019. Nat. Hazards. 2021, 108(3), 2721–38. DOI: 10.1007/s11069-021-04798-3.

3. Müller, M.; Dembélé S.; Zougmoré R.B.; Gaiser T.; Partey S.T. Performance of three sorghum cultivars under excessive rainfall and waterlogged conditions in the Sudano-Sahelian Zone of West Africa: A case study at the climate-smart village of Cinzana in Mali. Water. 2020, 72(10), 2655. DOI: 10.3390/w12102655

4. Alam, A.S.A.F.; Begum, H.; Masud, M.M.; Al-Amin, A.Q.; Filho, W.L. Agriculture insurance for disaster risk reduction: A case study of Malaysia. Int. J. Disaster Risk Reduct. 2020, 47, 9. DOI: 10.1016/j.ijdrr.2020.101626.

5. NOAA National Centers for Environmental Information (NCEI) U.S. billion-dollar weather and climate disasters. 2022. Available from: https://www.ncei.noaa.gov/access/billions/. DOI: 10.25921/stkw-7w73

6. Phukan, U.J.; Jeena, G.S.; Tripathi, V.; Shukla, R.K. *MaRAP2-4,* a waterlogging-responsive ERF from Mentha, regulates bidirectional sugar transporter *AtSWEETlŨ* to modulate stress response in *Arabidopsis*. Plant Biotechnol. J. 2018, 16(1), 221–33. DOI: 10.1111/pbi.12762.

7. Sasidharan, R.; Hartman, S.; Liu, Z.; Martopawiro, S.; Sajeev, N.; van Veen, H.; et al. Signal dynamics and interactions during flooding stress. Plant Physiol. 2018, 176(2), 1106–17. DOI: 10.1104/pp.17.01232.

8. Paul, P.L.C.; Bell, R.W.; Barrett-Lennard, E.G.; Kabir, E.; Mainuddin, M.; Sarker, K.K. Short-term waterlogging depresses early growth of sunflower *(Helianthus annuus* L.) on saline soils with a shallow water table in the coastal zone of Bangladesh. Soil Systems. 2021, 5(4), 12. DOI: 10.3390/soilsystems5040068.

9. Ide, R.; Ichiki, A.; Suzuki, T.; Jitsuyama, Y. Analysis of yield reduction factors in processing tomatoes under waterlogging conditions. Sci. Hortic. 2022, 295, 14. DOI: 10.1016/j.scienta.2021.110840.

10. Kaur, G.; Zurweller, B.A.; Nelson, K.A.; Motavalli, P.P.; Dudenhoeffer, C.J. Soil waterlogging and nitrogen fertilizer management effects on corn and soybean yields. Agron. J. 2017, 109(1), 97–106. DOI: 10.2134/agronj2016.07.0411

11. Lau, S.E.; Hamdan, M.F.; Pua, T.L.; Saidi, N.B.; Tan, B.C. Plant nitric oxide signaling under drought stress. Plants. 2021, 10(2), 29. DOI: 10.3390/plants10020360.

12. Wang, J.; Lv X.L.; Wang, H.W.; Zhong, X.M.; Shi, Z.S.; Zhu, M.; et al. Morpho-anatomical and physiological characteristics responses of a paried near-isogenic lines of waxy corn to waterlogging. Emir. J. FoodAgric. 2019, 31(12), 951–7. DOI: 10.9755/ejfa.2019.v31.i12.2045.

13. Pedersen, O.; Nakayama, Y.; Yasue, H.; Kurokawa, Y.; Takahashi, H.; Floytrup, A.H.; et al. Lateral roots, in addition to adventitious roots, form a barrier to radial oxygen loss in *Zea nicaraguensis* and a chromosome segment introgression line in maize. New Phytol. 2021, 229(1), 94–105. DOI: 10.1111/nph.16452.

14. Luan, H.; Guo, B.; Pan, Y.; Lv, C.; Shen, H.; Xu, R. Morpho-anatomical and physiological responses to waterlogging stress in different barley *(Hordeum vulgare* L.) genotypes. Plant Growth Regul. 2018, 85(3), 399–409. DOI: 10.1007/s10725-018-0401-9.

15. Amnan, M.A.M.; Pua, T.L.; Lau, S.E.; Tan, B.C.; Yamaguchi, H.; Hitachi, K.; et al. Osmotic stress in banana is relieved by exogenous nitric oxide. PeerJ. 2021, 9, 26. DOI: 10.7717/peerj.10879.

16. Pan, R.; Jiang, W.; Wang, Q.; Xu, L.; Shabala, S.; Zhang W.Y. Differential response of growth and photosynthesis in diverse cotton genotypes under hypoxia stress. Photosynthetica. 2019, 57(3), 772–9. DOI: 10.32615/ps.2019.087.

17. Qi, X.H.; Li, Q.Q.; Ma, X.T.; Qian, C.L.; Wang, H.H.; Ren, N.N.; et al. Waterlogging-induced adventitious root formation in cucumber is regulated by ethylene and auxin through reactive oxygen species signalling. Plant Cell Environ. 2019, 42(5), 1458–70. DOI: 10.1111/pce.13504.

18. Pan, J.W.; Sharif, R.; Xu, X.W.; Chen, X.H. Mechanisms of waterlogging tolerance in plants: Research progress and prospects. Front. Plant Sci. 2021, 11. DOI: 16, 10.3389/fpls.2020.627331.

19. Qi, X.; Li, Q.; Ma, X.; Qian, C.; Wang, H.; Ren, N.; et al. Waterlogging-induced adventitious root formation in cucumber is regulated by ethylene and auxin through reactive oxygen species signalling. Plant Cell Environ. 2019, 42(5), 1458–70. DOI: 10.1111/pce.13504

20. Liang, K.; Tang, K.; Fang, T.; Qiu, F. Waterlogging tolerance in maize: genetic and molecular basis. Mol. Breed. 2020, 40(12), 1–13. DOI: 10.1007/s11032-020-01190-0

21. Chin, W.Y.W.; Annuar, M.S.M.; Tan, B.C.; Khalid, N. Evaluation of a laboratory scale conventional shake flask and a bioreactor on cell growth and regeneration of banana cell suspension cultures. Sci. Hortic. 2014, 172, 39–46. DOI: 10.1016/j.scienta.2014.03.042.

22. FAOSTAT Statistical database. 2021. Available from: https://www.fao.org/faostat/en/#home.

23. Varma, V.; Bebber, D.P. Climate change impacts on banana yields around the world. Nat. Clim. Chang. 2019, 9(10), 752–7. DOI: 10.1038/s41558-019-0559-9.

24. Kanehisa, M.; Sato, Y.; Kawashima, M.; Furumichi, M.; Tanabe, M. KEGG as a reference resource for gene and protein annotation. Nucleic Acids Res. 2016, 44(D1), D457–62. DOI: 10.1093/nar/gkv1070.

25. Tavares, E.Q.P.; Grandis, A.; Lembke, C.G.; Souza, G.M.; Purgatto, E.; De Souza, A.P.; et al. Roles of auxin and ethylene in aerenchyma formation in sugarcane roots. Plant Signal. Behav. 2018, 13(3), 3. DOI: 10.1080/15592324.2017.1422464.

26. Yamauchi, T.; Tanaka, A.; Tsutsumi, N.; Inukai, Y.; Nakazono, M. A role for auxin in ethylene-dependent inducible aerenchyma formation in rice roots. Plants 2020, 9(5), 11. DOI: 10.3390/plants9050610.

27. Ploschuk, R.A.; Miralles, D.J.; Colmer, T.D.; Ploschuk, E.L.; Striker, G.G. Waterlogging of winter crops at early and late stages: impacts on leaf physiology, growth and yield. Front. Plant Sci. 2018, 9, 1863. DOI: 10.3389/fpls.2018.01863

28. Mhimdi, M.; Perez-Perez, J.M. Understanding of adventitious root formation: What can we learn from comparative genetics? Front. Plant Sci. 2020, 11, 10. DOI: 10.3389/fpls.2020.582020.

29. Mahmood, U.; Hussain, S.; Hussain, S.; Ali, B.; Ashraf, U.; Zamir, S.; et al. Morpho-physio-biochemical and molecular responses of maize hybrids to salinity and waterlogging during stress and recovery phase. Plants 2021, 10(7), 19. DOI: 10.3390/plants10071345.

30. Men, S.; Chen, H.; Chen, S.; Zheng, S.; Shen, X.; Wang, C.; et al. Effects of supplemental nitrogen application on physiological characteristics, dry matter and nitrogen accumulation of winter rapeseed *(Brassica napus* L.) under waterlogging stress. Sci. Rep. 2020, 10(1), 10201. DOI: 10.1038/s41598-020-67260-7.

31. Amnan, M.A.M.; Aizat, W.M.; Khaidizar, F.D.; Tan, B.C. Drought stress induces morpho-physiological and proteome changes of *Pandanus amaryllifolius*. Plants 2022, 11(2), 21. DOI: 10.3390/plants11020221.

32. Jia, L.T.; Qin, X.; Lyu, D.G.; Qin, S.J.; Zhang, P. ROS production and scavenging in three cherry rootstocks under short-term waterlogging conditions. Sci. Hortic. 2019, 257, 7. DOI: 10.1016/j.scienta.2019.108647.

33. Barickman, T.C.; Simpson, C.R.; Sams, C.E. Waterlogging causes early modification in the physiological performance, carotenoids, chlorophylls, proline, and soluble sugars of cucumber plants. Plants 2019, 8(6), 15. DOI: 10.3390/plants8060160.

34. Xiao, Y.S.; Wu, X.L.; Sun, M.X.; Peng, F.T. Hydrogen sulfide alleviates waterlogging-induced damage in peach seedlings via enhancing antioxidative system and inhibiting ethylene synthesis. Front. Plant Sci. 2020, 11, 14. DOI: 10.3389/fpls.2020.00696.

35. Xie, R.J.; Zheng, L.; Jiao, Y.; Huang, X. Understanding physiological and molecular mechanisms of citrus rootstock seedlings in response to root zone hypoxia by RNA-Seq. Environ. Exp. Bot. 2021, 192, 14. DOI: 10.1016/j.envexpbot.2021.104647.

36. Da-Silva, C.J.; do Amarante, L. Time-course biochemical analyses of soybean plants during waterlogging and reoxygenation. Environ. Exp. Bot. 2020, 180, 11. DOI: 10.1016/j.envexpbot.2020.104242.

37. Li, H.; Wang, H.; Wen, W.J.; Yang, G.W. The antioxidant system in *Suaeda salsaunder* salt stress. Plant Signal. Behav. 2020, 15(7), 6. DOI: 10.1080/15592324.2020.1771939.

38. Anee, T.I.; Nahar, K.; Rahman, A.; Al Mahmud, J.; Bhuiyan, T.F.; Ul Alam, M.; et al. Oxidative damage and antioxidant defense in *Sesamum indicum* after different waterlogging durations. Plants 2019, 5(7), 18. DOI: 10.3390/plants8070196.

39. Azahar, I.; Ghosh, S.; Adhikari, A.; Adhikari, S.; Roy, D.; Shaw, A.K.; et al. Comparative analysis of maize root sRNA transcriptome unveils the regulatory roles of miRNAs in submergence stress response mechanism. Environ. Exp. Bot. 2020, 171, 20. DOI: 10.1016/j.envexpbot.2019.103924.

40. Khan, M.; Trivellini, A.; Chhillar, H.; Chopra, P.; Ferrante, A.; Khan, N.A.; et al. The significance and functions of ethylene in flooding stress tolerance in plants. Environ. Exp. Bot. 2020, 179, 13. DOI: 10.1016/j.envexpbot.2020.104188.

41. Perata, P. Ethylene signaling controls fast oxygen sensing in plants. Trends Plant Sci. 2020, 25(1), 3–6. DOI: 10.1016/j.tplants.2019.10.010.

42. Loreti, E.; Poggi, A.; Novi, G.; Alpi, A.; Perata, P. A genome-wide analysis of the effects of sucrose on gene expression in Arabidopsis seedlings under anoxia. Plant Physiol. 2005, 137(3), 1130–8. DOI: 10.1104/pp.104.057299.

43. Hartman, S.; van Dongen, N.; Renneberg, D.; Welschen-Evertman, R.A.M.; Kociemba, J.; Sasidharan, R.; et al. Ethylene differentially modulates hypoxia responses and tolerance across *Solanum* species. Plants 2020, 9(8), 14. DOI: 10.3390/plants9081022.

44. Tsuchiya, Y.; Nakamura, T.; Izumi, Y.; Okazaki, K.; Shinano, T.; Kubo, Y.; et al. Physiological role of aerobic fermentation constitutively expressed in an aluminum-tolerant cell line of tobacco *(Nicotiana tabacum)*. Plant Cell Phys. 2021, 62(9), 1460–77. DOI: 10.1093/pcp/pcab098.

45. Licausi, F.; van Dongen, J.T.; Giuntoli, B.; Novi, G.; Santaniello, A.; Geigenberger, P.; et al. *HRE1* and *HRE2*, two hypoxia-inducible ethylene response factors, affect anaerobic responses in *Arabidopsis thaliana*. Plant J. 2010, 62(2), 302–15. DOI: 10.1111/j.1365-313X.2010.04149.x.

46. Wei, X.; Xu, H.; Rong, W.; Ye, X.; Zhang, Z. Constitutive expression of a stabilized transcription factor group VII ethylene response factor enhances waterlogging tolerance in wheat without penalizing grain yield. Plant Cell Environ. 2019, 42(5), 1471–85. DOI: 10.1111/pce.13505

47. Tamang, B.G.; Li, S.; Rajasundaram, D.; Lamichhane, S.; Fukao, T. Overlapping and stress-specific transcriptomic and hormonal responses to flooding and drought in soybean. Plant J. 2021, 107(1), 100-17. DOI: 10.1111/tpj.15276.

48. Harkey, A.F.; Watkins, J.M.; Olex, A.L.; DiNapoli, K.T.; Lewis, D.R.; Fetrow, J.S.; et al. Identification of transcriptional and receptor networks that control root responses to ethylene. Plant Physiol. 2018, 176(3), 2095–118. DOI: 10.1104/pp.17.00907.

49. Dawood, T.; Yang, X.P.; Visser, E.J.W.; Te Beek, T.A.H.; Kensche, P.R.; Cristescu S.M.; et al. A coopted hormonal cascade activates dormant adventitious root primordia upon flooding in *Solanum dulcamara*. Plant Physiol. 2016, 170(4), 2351–64. DOI: 10.1104/pp.15.00773.

50. Nguyen, T-N.; Tuan, P.A.; Mukherjee, S.; Son, S.; Ayele, B.T. Hormonal regulation in adventitious roots and during their emergence under waterlogged conditions in wheat. J. Exp. Bot. 2018, 69(16), 4065–82. DOI: 10.1093/jxb/ery190

51. Shimamura, S.; Yoshioka, T.; Yamamoto, R.; Hiraga, S.; Nakamura, T.; Shimada, S.; et al. Role of abscisic acid in flood-induced secondary aerenchyma formation in soybean (*Glycine max*) hypocotyls. Plant Prod. Sci. 2014, 17(2), 131–7. DOI: 10.1626/pps.17.131.

52. Khan, M.I.R.; Trivellini, A.; Chhillar, H.; Chopra, P.; Ferrante, A.; Khan, N.A.; et al. The significance and functions of ethylene in flooding stress tolerance in plants. Environ. Exp. Bot. 2020, 104188. DOI: 10.1016/j.envexpbot.2020.104188

53. Bashar, K.K.; Tareq, M.Z.; Islam, M.S. Unlocking the mystery of plants’ survival capability under waterlogging stress. PlantSci. Today. 2020, 7(2), 142–53. DOI: 10.14719/pst.2020.7.2.663.

54. Zhang, P.; Lyu D.; Jia, L.; He, J.; Qin, S. Physiological and *de novo* transcriptome analysis of the fermentation mechanism of *Cerasus sachalinensis* roots in response to short-term waterlogging. BMC Genomics. 2017, 18(1), 649. DOI: 10.1186/s12864-017-4055-1.

55. Borrego-Benjumea, A.; Carter, A.; Tucker, J.R.; Yao, Z.; Xu, W.; Badea, A. Genome-wide analysis of gene expression provides new insights into waterlogging responses in barley *(Hordeum vulgare* L.). Plants 2020, 9(2), 23. DOI: 10.3390/plants9020240.

56. Ren, B.Z.; Dong, S.T.; Zhao, B.; Liu, P.; Zhang, J.W. Responses of nitrogen metabolism, uptake and translocation of maize to waterlogging at different growth stages. Front. Plant Sci. 2017, 8, 9. DOI: 10.3389/fpls.2017.01216.

57. Kamsen, R.; Kalapanulak, S.; Chiewchankaset, P.; Saithong, T. Transcriptome integrated metabolic modeling of carbon assimilation underlying storage root development in cassava. Sci. Rep. 2021, 11(1), 15. DOI: 10.1038/s41598-021-88129-3.

58. Turner, N.C. Techniques and experimental approaches for the measurement of plant water status. Plant Soil 1981, 58(1-3), 339-66, DOI: 10.1007/bf02180062.

59. Mano, Y.; Omori, F. High-density linkage map around the root aerenchyma locus *Qaer1. 06* in the backcross populations of maize Mi29× teosinte *“Zea nicaraguensis*”. Breed. Sci. 2009, 59(4), 427–33. DOI: 10.1270/jsbbs.59.427

60. Junglee, S.; Urban L.; Sallanon H.; Lopez-Lauri F.J.A.Jo.A.C. Optimized assay for hydrogen peroxide determination in plant tissue using potassium iodide. Am. J. Anal. Chem. 2014, 5(11), 730. DOI: 10.4236/ajac.2014.511081

61. Bates, L.S.; Waldren, R.P.; Teare, I. Rapid determination of free proline for water-stress studies. Plant Soil 1973, 39(1), 205–7. DOI: 10.1007/BF00018060

62. Zhang, Q.; Zhang, J.Z.; Chow, W.S.; Sun, L.L.; Chen, J.W.; Chen, Y.J.; et al. The influence of low temperature on photosynthesis and antioxidant enzymes in sensitive banana and tolerant plantain *(Musa* sp.) cultivars. Photosynthetica. 2011, 49(2), 201–8. DOI: 10.1007/s11099-011-0012-4.

63. Carlberg, I.; Mannervik, B. Purification and characterization of the flavoenzyme glutathione reductase from rat liver. J. Bio.l Chem. 1975, 250(14), 5475–80. DOI: 10.1016/S0021-9258(19)41206-4

64. Pütter, J. Peroxidases. Methods of enzymatic analysis: Elsevier, 1974, 685–90. DOI: 10.1016/B978-0-12-091302-2.50033-5

65. Asif, M.H.; Dhawan, P.; Nath, P. A simple procedure for the isolation of high quality RNA from ripening banana fruit. Plant Mol. Biol. Rep. 2000, 18(2), 109–15. DOI: 10.1007/BF02824018

66. Pfaffl, M.W. A new mathematical model for relative quantification in real-time RT-PCR. Nucleic Acids Res. 2001, 29(9), e45. DOI: 10.1093/nar/29.9.e45.

67. FastQC: A quality control tool for high throughput sequence data. 2010. Available from: http://www.bioinformatics.babraham.ac.uk/projects/fastqc/.

68. Martin, M. Cutadapt removes adapter sequences from high-throughput sequencing reads. EMBnetjournal. 2011. DOI: 10.14806/ej.17.1.200.

69. Droc, G.; Lariviere, D.; Guignon, V.; Yahiaoui, N.; This, D.; Garsmeur, O.; et al. The Banana Genome Hub. Database. 2013, 14. DOI: 10.1093/database/bat035.

70. Kim, D.; Paggi, J.M.; Park, C.; Bennett, C.; Salzberg, S.L. Graph-based genome alignment and genotyping with HISAT2 and HISAT-genotype. Nat. Biotechnol. 2019, 37(8), 907. DOI: 10.1038/s41587-019-0201-4.

71. Love, M.I.; Huber, W.; Anders, S. Moderated estimation of fold change and dispersion for RNA-seq data with DESeq2. Genome Biol. 2014, 15(12), 38. DOI: 10.1186/s13059-014-0550-8.

72. Young, M.D.; Wakefield M.J.; Smyth G.K.; Oshlack A. Gene ontology analysis for RNA-seq: accounting for selection bias. Genome Biol. 2010, 11(2). DOI: 10.1186/gb-2010-11-2-r14.

