## Supplementary figures and images for "Waterlogging stress induces antioxidant defense responses and aerenchyma formation and alters metabolisms of banana plants"

### Supplemental Figure 1

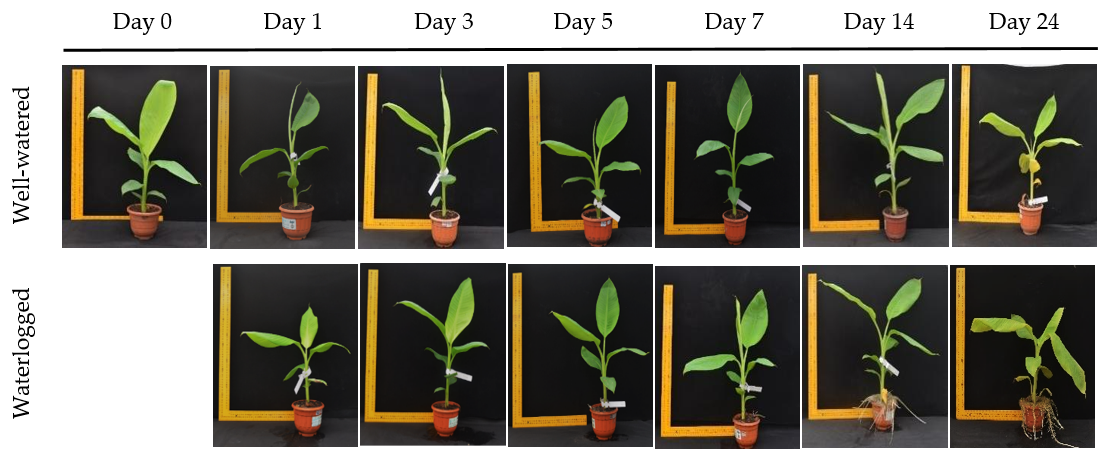
